# Joint-specific rheumatoid arthritis fibroblast-like synoviocyte regulation identified by integration of chromatin access and transcriptional activity

**DOI:** 10.1101/2024.01.12.575379

**Authors:** Eunice Choi, Camilla R. L. Machado, Takaichi Okano, David Boyle, Wei Wang, Gary S. Firestein

## Abstract

The mechanisms responsible for the distribution and severity of joint involvement in rheumatoid arthritis (RA) are not known. To explore whether site-specific FLS biology might be associated with location-specific synovitis and explain the predilection for hand (wrist/metacarpal phalangeal joints) involvement in RA, we generated transcriptomic and chromatin accessibility data from FLS to identify the transcription factors (TFs) and pathways. Networks were constructed by integration of chromatin accessibility and gene expression data. Analysis revealed joint-specific patterns of FLS phenotype, with proliferative, migratory, proinflammatory, and matrix-degrading characteristics observed in resting FLS derived from the hand joints compared with hip or knee. TNF-stimulation amplified these differences, with greater enrichment of proinflammatory and proliferative genes in hand FLS compared with hip and knee FLS. Hand FLS also had the greatest expression of markers associated with an ‘activated’ state relative to the ‘resting’ state, with the greatest cytokine and MMP expression in TNF-stimulated hand FLS. Predicted differences in proliferation and migration were biologically validated with hand FLS exhibiting greater migration and cell growth than hip or knee FLS. Distinctive joint-specific FLS biology associated with a more aggressive inflammatory response might contribute to the distribution and severity of joint involvement in RA.

## Introduction

Rheumatoid arthritis (RA) is a common form of inflammatory arthritis characterized by autoimmunity and symmetric, polyarticular synovitis^1^. RA fibroblast-like synoviocytes (FLS) are key effector cells in the synovium that are imprinted by their environment and display an aggressive phenotype^2,3^. For example, rheumatoid FLS produce a variety of proteases that damage the extracellular matrix and enable invasion of synovial tissue into bone and cartilage^2^. These cells also produce cytokines such as IL-6^4^ and GM-CSF that activate nearby immune effector cells like B and T lymphocytes^5,6^. Targeting FLS has been proposed as a strategy to avoid typical side effects like systemic immune suppression associated with many current RA therapies^2^.

RA is characterized by distinct anatomic and temporal patterns of joint involvement. The small joints of the hands and feet are often affected in very early RA; although larger joints like hips and knees can be affected early, joint damage in these locations typically occurs later in the course of disease. Biomechanics^7–9^, lymphatic distribution^10^ and innervation^11^ might contribute to the joint distribution in RA, but it is possible that location-specific FLS biology contributes^12–14^. For example, we previously showed that FLS isolated from different joints have disease-independent and disease-specific methylation and transcriptomic patterns that are related to joint development and inflammatory pathways, respectively^14^. Others have also shown that FLS from various joints have distinct characteristics and differential responses to the proinflammatory microenvironment^12,13^. Some of which might be related to increased expression of the lncRNA HOTAIR in knee FLS, although this only contributes to about half of the differences with hand compared with knee and hip FLS^12^. These data suggest that disease processes might vary from joint to joint and that the full extent of joint-specific regulatory networks needs to be defined.

To extend this work, RNA-seq and ATAC-seq of resting and TNF-stimulated FLS derived from 3 anatomical location in patients with RA were conducted to evaluate the FLS states in the stable and inflammatory contexts. We demonstrate that FLS from hand, knee and hip have distinct transcriptomes and chromatin accessibility patterns. By integrating transcriptomes and epigenomes using our novel Taiji method^15,16^, we identified key transcription factor drivers that translate into unique joint-specific phenotypes, including predicted drivers that increase HOTAIR expression specifically in hand FLS. Moreover, joint-specific TFs and regulatee chromatin accessibility and expression patterns correlate with patterns of joint involvement and severity in RA, namely with the greatest proinflammatory, proliferative, and migratory phenotype observed in hand FLS. Computational predictions were then biologically validated. These findings might contribute to the typical cadence of disease onset and progression in RA.

## Results

### Computational and experimental overview

To test the hypothesis that fibroblast-like synoviocytes (FLS) have distinct joint-specific phenotypes and behavior, we characterized FLS transcriptomes and chromatin accessibility in RA FLS obtained from synovial tissue isolated from 3 different joints (10 hand (metacarpal phalangeal (MCP) joints/wrists), 10 hips, and 10 knees). These cells were studied in their basal condition as well as after TNF stimulation. Figure 1A shows the workflow for this process (see also computational methods).

**Figure 1.**
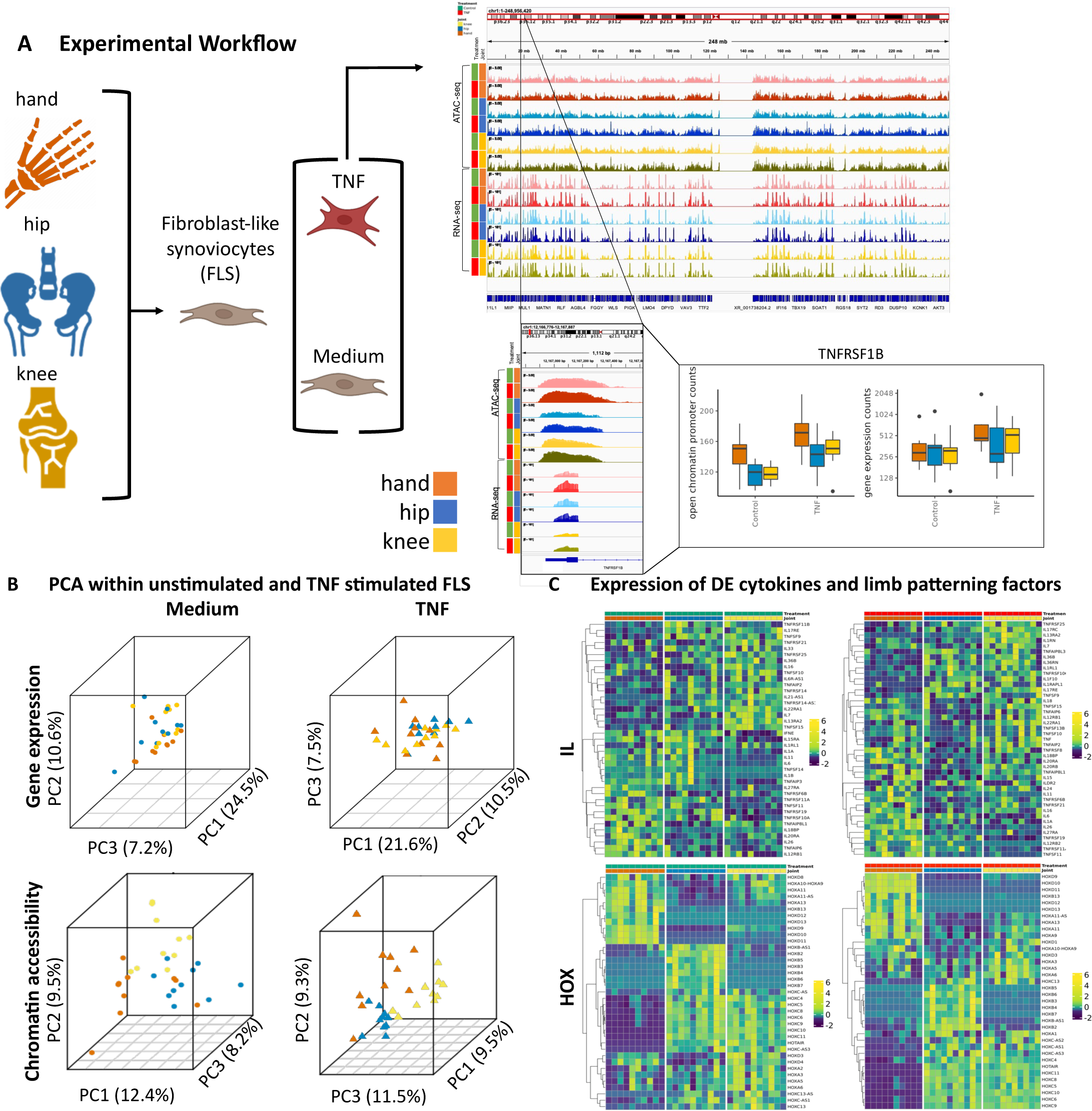
**A.** Experimental workflow. Synovial tissue was obtained from RA patients at the time of clinically indicated arthroplasty at 3 joint locations: knee, hip and hand (metacarpal phalangeal joint and wrist). Cells were cultured in medium or TNF-stimulated conditions and harvested for whole genome RNA-seq and ATAC-seq. **Amber, blue, and yellow** represent hand, hip, and knee, respectively. **Green and red** represent medium and TNF-stimulated conditions, respectively. The color palette is maintained throughout all figures. **B.** PCA of all chromatin accessibility and gene expression profiles within medium and TNF-stimulated conditions. PCA using the gene expression profiles within both medium and TNF- stimulated conditions revealed the greatest differences between hand and hip. PCA using the chromatin accessibility profiles revealed greater stratification between the joints compared to gene expression. In medium, all joints were separated. In TNF-stimulated conditions, the segregation between hand vs hip and knee were amplified. **C.** Gene expression profiles of differentially expressed interleukins and homeobox (HOX) genes. RA FLS have distinct cytokine expression in medium. Similarly, RA FLS have distinct patterns of cytokine induction after TNF stimulation. Several proinflammatory cytokines like IL-6 had marked induction in hand in response to TNF (FC between TNF-stimulated hand / knee: 2.0, hand /hip: 1.12, hip / knee: 1.7). Limb patterning HOX gene expression were evaluated for medium and TNF- stimulated conditions. Hand FLS had the most distinct expression compared to hip and knee within both medium and TNF-stimulated conditions for several HOX features like HOXD10. See supplementary file 3 for DEGs, p-values, and FCs.

#### Stratifying RA FLS gene expression based on joint location

Joint-specific transcriptional patterns were identified by first focusing our analysis on unstimulated cultured FLS. Principal component analysis (PCA) was performed on all gene expression profiles to summarize global patterns of gene expression and observed joint-specific differences, with the greatest separation between hand and hip (Figure 1B). We then evaluated the PCA of gene expression profiles for TNF-stimulated FLS. As with unstimulated FLS, the greatest differences were noted between hand and hip while knee was intermediate (Figure 1B). Gene expression patterns in PC space between medium and TNF-stimulated FLS from the same joint location were then examined. The PCA revealed that joint-specific transcriptome differences were amplified by TNF, as shown by greater variance represented by PC1 in TNF-stimulated cell lines. Supplementary files 1, 2, and 3 show the differentially expressed genes and their respective pathways between joint locations (hand-knee; hand-hip; and knee-hip) for unstimulated and TNF-stimulated FLS.

Because cytokines participate in the pathogenesis of RA^1,17^, we focused our next analysis on cytokine expression patterns between joint locations for unstimulated and TNF-stimulated FLS. We have previously shown that IL-6 is differentially expressed and methylated between hip and knee FLS^14,18^. First, we evaluated joint-specific expression within unstimulated FLS (Figure 1C). For cytokines that were differentially expressed in at least one pairwise comparison, we determined joint-specific expression patterns. FLS exhibited joint-specific expression of proinflammatory cytokines with the greatest expression of cytokines observed in hand (16 cytokines), knee as intermediate (13 cytokines) and the least in hip (10 cytokines) (Figure 1C). As a supplementary analysis, we performed hierarchical clustering using Ward’s Hierarchical Agglomerative Clustering Method (ward.d2) to determine expression patterns across joints.

Unstimulated FLS exhibited joint-specific stratification with the greatest mixing between hip and knee FLS (Supplementary figure 1). We then evaluated joint-specific expression within TNF-stimulated FLS. After TNF stimulation, the pattern persisted with the greatest expression of differentially expressed cytokines observed in hand (21 cytokines), knee as intermediate (13 cytokines), and the least in hip (5 cytokines). Again, we performed hierarchical clustering and found that TNF-stimulated hand FLS were clearly segregated from hip and knee FLS. This reflects hand-specific cytokine response to TNF and is reflected by marked induction of several proinflammatory cytokines like IL-6 and IL1A enriched in hand (Figure 1C, Supplementary figure 1). These data suggest that RA FLS have distinct cytokine expression and responses to TNF based on joint location and might reflect RA severity.

We then corroborated joint-specific expression of limb patterning genes as they may contribute to defining joint-specific function and biology across RA patients^14^. We evaluated expression of HOX genes between the FLS derived from hip, knee and hand joints (Figure 1C). First, we evaluated joint-specific expression within unstimulated FLS. For limb patterning genes that were differentially expressed in at least one pairwise comparison within unstimulated FLS, we determined joint-specific expression patterns. As previously noted, FLS exhibited distinct joint-specific expression of several HOX genes^12–14^, including HOXD10 (Lee, H et al, submitted for publication) (Figure 1C). The lncRNA HOTAIR was previously implicated as a regulator that contributes to these differences^13^. Indeed, we confirmed that HOTAIR expression is greater in knee and hip FLS compared with hand FLS in unstimulated and TNF stimulated conditions (Supplementary figure 2, Supplementary figure 4). Our data suggests that these differences are likely related to decreased chromatin accessibility of HOTAIR regulatory regions in unstimulated hand FLS (Supplementary figure 2, Supplementary figure 3).

Overall gene expression patterns were also evident under TNF-stimulated conditions, with consistently decreased expression in hand compared to hip or knee FLS (Figure 1C). Again, we performed hierarchical clustering focused on HOX genes. After TNF stimulation, the topologically distinct patterns of HOX gene expression were still with greater similarity between hip with knee compared with hand FLS (Supplementary figure 5).

#### Stratifying RA FLS chromatin accessibility based on joint location

Previous studies determined distinct epigenetic patterns that reflect anatomic location, including differential methylation^14,19^, histone marks^20^, lncRNA^13^, and locus accessibility^20^ signatures. We conducted PCA on chromatin accessibility profiles with unstimulated FLS data (Figure 1B). PCA revealed site-specific stratification that was more prominent than the transcriptome. We then evaluated the PCA of chromatin accessibility profiles of TNF stimulated FLS. (Figure 1B). All joint locations segregated; like unstimulated FLS, TNF-stimulated FLS from hand had the greatest differences and separation from knee and hip and amplified the differences observed in unstimulated cells. Supplementary files 1 and 4 show differential chromatin accessibility and the respective pathways between joint locations (hand-knee; hand-hip; and knee-hip) for unstimulated and TNF-stimulated FLS.

### Joint-specific patterns of unstimulated RA FLS transcriptional regulation

#### Unstimulated FLS have joint-specific transcriptional regulation

Separate analysis of individual datasets like chromatin accessibility and the transcriptome provides some information on the functional differences in FLS based on joint location. However, to understand the distinct joint-specific networks in greater depth, ATAC-seq and RNA-seq data were integrated using our novel Taiji^15,16^ pipeline (see Methods). Taiji constructs networks using gene expression and open chromatin profiles and applies the PageRank score on all regulator nodes to quantify TF importance (Methods). Using this method, we identified shared and joint-specific distinct TF PageRank scores in unstimulated FLS (Figure 2A). Using pairwise comparisons between joints, we identified 73, 115, and 140 differentially ranked TFs (Wilcoxon test, p-value < 0.05) between hand vs knee, hand vs hip, and knee vs hip, respectively (Figure 2A, 2B) (Supplementary file 5). Hand FLS had increased PageRanks for TFs implicated in immune responses and inflammation, such as cytokine signaling, Toll-like receptor (TLR), and mitogen activated protein kinase (MAPK) cascades (e.g., SMAD3, ATF1, ATF2, CREB1) (Figure 2C). Knee FLS were notable for the increased importance of TFs involved in interferon and NOTCH signaling (e.g., IRF2, STAT1, STAT2, HEY1, PBX1). Hip FLS, on the other hand, had increased importance of TFs focused on pathways like RUNX1 signaling (ELF1, LMO2, MYB) (Supplementary file 5). These data indicate that FLS from the different joints have distinct regulators that drive differential FLS function and behavior.

**Figure 2.**
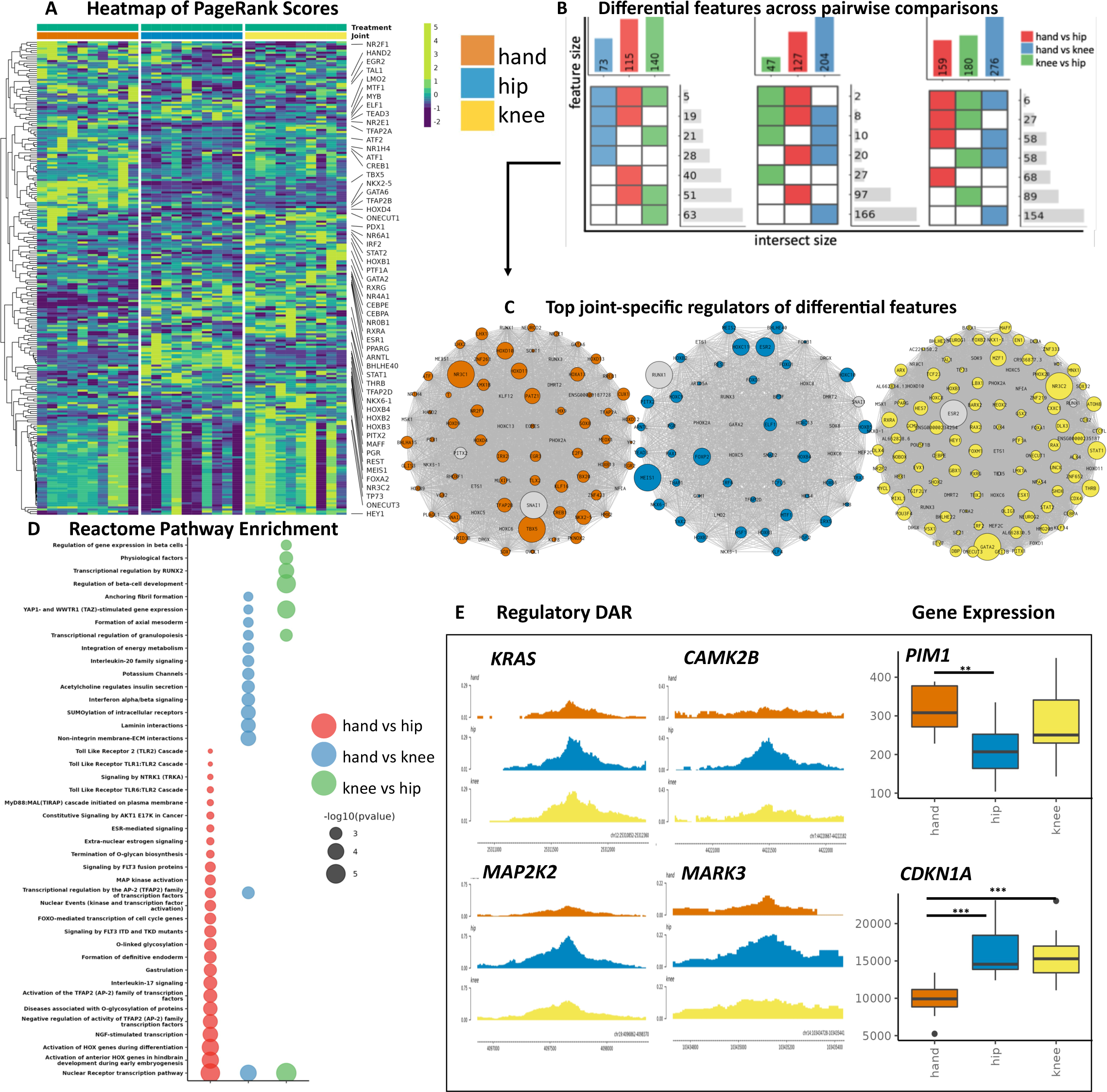
**A**. Heatmap of PageRanks of the top differential TFs selected from pairwise comparisons between unstimulated FLS from hand, hip, and knee. Pathway enriched TFs are labeled. For example, CAMP responsive element binding protein 1 (CREB1) has greatest importance in hand FLS. Important TFs are reflected by increased accessibility of loci containing the TF motif, increased TF expression, and increased predicted TF binding affinity of the identified motif within the open chromatin regions. **B**. Intersection plot of differential regulators (TFs) and regulatees (DAR and DEGs that are regulated by the differential regulator). Top bars indicate all differential features for each pairwise comparison. Heatmap indicate shared differential comparisons. Side bars indicate number of features that are shared between comparisons. For example, hand vs hip has 115 TFs that were identified with differential PageRanks. 51 of those differential TFs were also identified from hip vs knee. **Blue, red, and green** are used for hand vs knee, hand vs hip, and knee vs hip comparisons, respectively. **C**. For each joint, differential TFs with the greatest PageRank score in that joint was visualized. Nodes are scaled by PageRank. For example, PATZ1 has differential PageRanks between hand vs hip and has the greatest PageRank in hand (FC in PATZ1 PageRank for hand vs knee: 1.2, hand vs hip: 1.3, knee vs hip: 0.8). **D**. Pathway enrichment analysis of TFs and regulatee DARs and DEGs between pairwise comparisons. The greatest bias towards FLS aggressiveness was observed in hand vs hip comparison and least between hip vs knee comparisons. Hand vs hip and hand vs knee comparisons were enriched for proinflammatory *e.g. MAP kinase activation* (p-value < 0.003), and proliferative pathways *e.g. Constitutive Signaling by AKT1 E17K in Cancer* (p-value < 0.006). **E**. Visualization of selected features. Genome browser are differential open chromatin regulatory loci and boxplots are DEGs as representative examples of differential proliferative and proinflammatory features. P-values are **\*, **, \*\*\*** for p-value < 0.05, 0.01, 0.005, respectively.

#### Unstimulated FLS have joint-specific TF regulatee profiles

We then evaluated joint-specific regulatees of these differential TFs in unstimulated FLS. To infer regulatory mechanisms of FLS phenotype, regulatees analyzed focused on differentially expressed (DEGs) or accessible regions (DARs) (Figure 2B). Of interest, only limited overlap was observed between DEGs and DARs, likely because FLS were unstimulated and the regulatory regions are in an epigenetically poised rather than active state (Supplementary figure 6). Evaluation of DEGs and DARs revealed distinct joint-specific drivers (Figure 2C). Pathway enrichment analysis of the union of the differential features (Figure 2B) revealed a gradient in FLS functional differences, with the greatest bias towards FLS aggressiveness between hand compared to hip and least between hip compared to knee (Figure 2D). This gradient could reflect the pattern of disease onset severity in joints involved in RA, which is often affected early in metacarpal-phalangeal joints and wrists. Hand vs hip and hand vs knee comparisons were enriched for proinflammatory and proliferative pathways in hand FLS with minimal overlap for knee vs hip comparisons. Evaluation of DARs revealed FLS have joint-specific chromatin accessibility in key regulatory regions that confer well-characterized aggressive behaviors of RA FLS^2^ (Figure 2E). For example, hand FLS had decreased expression of cell-cycle inhibitors and increased expression of proto-oncogenes that are associated with cell growth (Figure 2E).

We also evaluated whether the potential regulator HOTAIR might contribute to joint-specific regulatory mechanisms in hand FLS. Due to high sample-specific heterogeneity in TF-HOTAIR edges, we applied a moderate filter of TF-HOTAIR edges that were identified in at least 20% of samples within each joint and treatment (unstimulated and TNF-stimulated). Taiji identified 0, 60, and 71 putative regulators of HOTAIR in unstimulated hand, hip, and knee, respectively, with 54 shared TFs between hip and knee (Supplementary figure 7). TFs were ranked by TF- HOTAIR edge weights which defines TF-gene regulatory potential to predict the most robust regulators that drive the hand FLS-specific phenotype. No predicted TFs for hand FLS were identified, which correlated with the lack of HOTAIR expression (Supplementary figure 2, Supplementary figure 3). However, hip and knee FLS had distinct predicted regulators and TF- HOTAIR edge weights (Supplementary figure 7). For example, hip and knee shared the top regulators (e.g., STAT1, STAT6, and MAZ), suggesting that similar regulatory mechanisms defined by epigenetics may drive increased HOTAIR expression in hip and knee FLS (Supplementary figure 7).

### TNF amplifies joint-specific differences in RA FLS transcriptional regulation

#### FLS have joint-specific responses in transcriptional regulation

Joint-specific responses in TF PageRanks were then evaluated by comparing unstimulated vs TNF-stimulated FLS within each joint. As expected, proinflammatory TFs like NFKB1, NFKB2 and RELB had significantly increased PageRanks after TNF stimulation for FLS derived from all joint locations (Figure 3A, Supplementary file 5) (Wilcoxon test, p-value < 0.05). Among the shared differential TFs, hand FLS compared to other joints had increased induction of proinflammatory TFs, such as *RELA, RELB* and *NFKB2*. Furthermore, hand FLS had unique differential TF PageRanks associated with proliferation, such as *E2F1, TFDP1 and FOXO4* and proinflammatory drivers, like *STAT3, RREB1* and *CEBPD* (Supplementary file 5). The observations suggest that TNF-stimulated hand FLS have epigenetic and transcriptomic responses that disproportionately enhance proinflammatory and proliferative TFs functions compared with knee or hip FLS.

**Figure 3.**
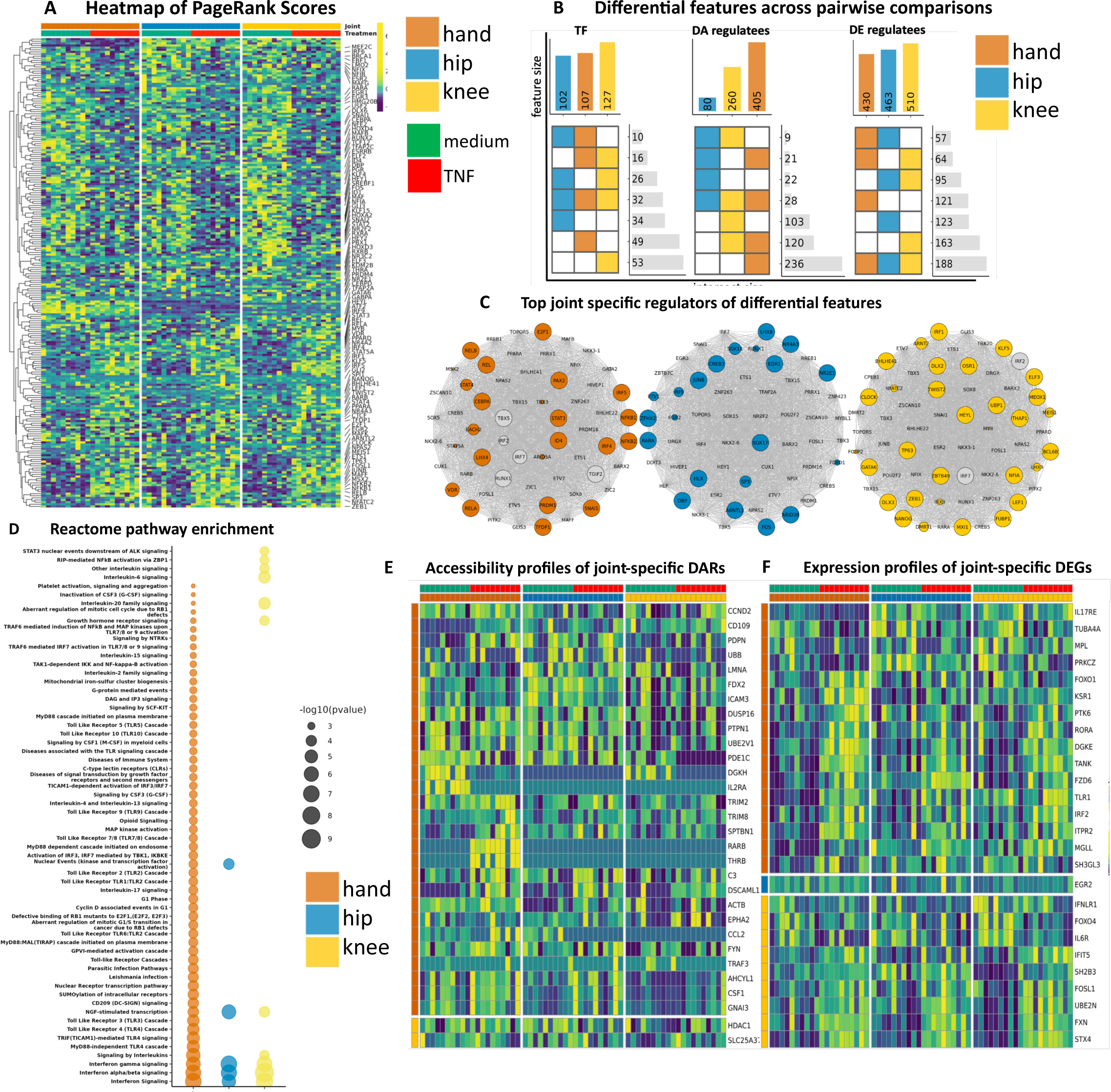
**A**. Heatmap of PageRanks of the top differential TFs selected from within joints and between unstimulated vs TNF stimulated FLS. Hand FLS had unique increased importance of proliferative *drivers i.e. E2F1, TFDP1 and FOXO4* and proinflammatory *drivers i.e. STAT3, RREB1 and CEBPD.* See supplementary file 5 for differential TFs, p-values, and FC within each joint and between unstimulated vs TNF-stimulated FLS.**B**. Intersection plot of differential regulators (TFs) and regulatees (DAR and DEGs that are regulated by the differential regulator). Top bars indicate all differential features for each pairwise comparison. Heatmap indicate shared differential comparisons. Side bars indicate number of features that are shared between comparisons. **C**. For each joint, differential TFs between unstimulated vs TNF stimulated FLS that regulate DEG/DARs are visualized. Nodes are ranked by FC i.e. large nodes indicate increased PageRank in TNF stimulated vs unstimulated for that joint. Colored nodes have greatest FC in that joint. Gray nodes have greater FC in another joint (although it is still differential for that particular joint). **D**. Pathway enrichment analysis of the union of TFs and regulatee DARs and DEGs between unstimulated vs TNF stimulated FLS. Hand FLS exhibits unique enrichment of several proinflammatory *e.g. MyD88-independent TLR4 cascade* (p-value < 1E-05) and proliferative e.g. *Aberrant regulation of mitotic cell cycle due to RB1 defects* (p-value < 0.002) pathways. **E**. Joint-specific DARs that are pathway enriched between unstimulated vs TNF stimulated FLS*, e.g.,* genes that are only differentially accessible in one joint. Sum of all differentially accessible regulatory loci are visualized. Hand FLS exhibits the most dynamic chromatin accessibility in response to TNF. **F**. Joint-specific DEGs that are pathway enriched between unstimulated vs TNF stimulated FLS. Hand FLS exhibit the most aggressive and joint-specific transcriptomic changes in response to stimuli.

#### TNF amplifies joint-specific differences in regulatee profiles

The regulatees of differential TFs were then evaluated after TNF stimulated, again focusing on DEGs and DARs (Figure 3B). TNF-stimulated FLS had joint-specific TFs and that reflected site-specific epigenetic and transcriptomic responses (Figure 3C). Pathway enrichment of the union of all differential features (Figure 3B) again revealed a gradient in proinflammatory responses, with the greatest and least proinflammatory pathways observed in hand and hip, respectively, with knee as intermediate (Figure 3D). For example, hand FLS had unique enrichment of proliferative pathways like *G1 Phase* and *Cyclin D associated events in G1* (Figure 3B, Supplementary file 2). Evaluation of joint-specific and pathway enriched DARs revealed hand FLS might also have significant epigenetic plasticity (Figure 3B, 3E) that correlates with hand-specific differential PageRanks in chromatin remodeling TFs, such as *CTCF, ATF2, KDM2B, SPI1 and TFAP2C* (Wilcoxon test, p-value < 0.05). Furthermore, evaluation of pathways enriched and their unique DEGs revealed that hand FLS have the greatest expression of proinflammatory and proliferative genes (Figure 3F). As with our analysis of unstimulated FLS, decreased HOTAIR chromatin accessibility was observed (Supplementary figure 1, Supplementary figure 2). We identified key regulators of HOTAIR after TNF-stimulation and identified 0, 22, and 36 putative regulators of HOTAIR for hand, hip, and knee, respectively, and identified the drivers in the latter two included pro-inflammatory TFs like NFKB1, RELB and IRF1 (Supplementary figure 7). This suggests a more comprehensive mechanism to explain joint-specific response to TNF and potentially clarifies the role of HOTAIR in the context of each transcriptional network to partially drive joint-specific FLS function. From the evaluation of joint-specific differences in PageRanks, hand-specific differences observed in unstimulated FLS were amplified by TNF and reflected by greater numbers of joint-specific pathways and transcriptomic and epigenomic responses compared with hip and knee FLS.

### Validation of computational predictions

#### Hand FLS have significant enrichment of genes associated with the ‘activated’ state

Recent studies of fibroblast states in RA synovium showed that these cells can exist in ‘activated’ and ‘resting’ states^21^. Given our prediction of increased pathogenicity of hand FLS, we tested whether hand FLS are enriched for genes associated with activated FLS states as defined by that report. Hand and knee had increased expression of activation state markers; however, knee FLS also had enrichment of resting markers (Figure 4A). To clarify the cell states for FLS from different joint locations, we evaluated the ratio of activated to resting genes for each joint. We found that hand FLS have a significantly increased ratio of activation markers compared with hip or knee FLS using either DEGs or all markers (Permutation test, p-value < 0.05) (Figure 4B, Supplementary figure 8, Supplementary figure 9). These data suggest that the RA hand synovium is enriched for FLS in the activated state and is consistent with our prediction that hand FLS are more likely to be aggressive.

**Figure 4.**
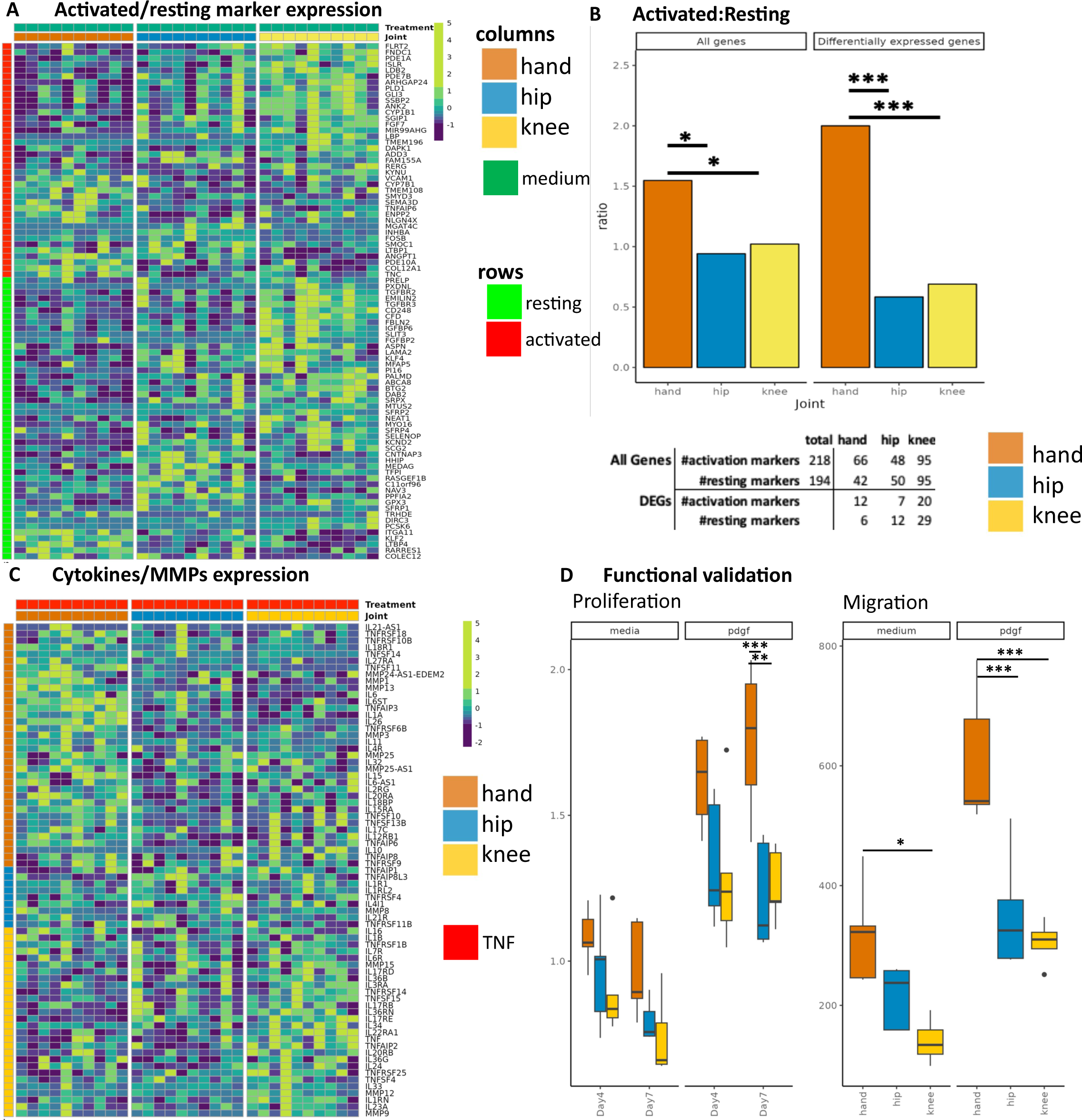
**A.** Heatmap of differentially expressed activated and resting marker expression in unstimulated FLS. Rows are activated/resting markers. While both hand and knee have increased expression of activated markers, knee also has increased expression of resting markers, indicating hand FLS has greatest ratio of activated to resting marker expression. **B**. Ratio of number of markers of maximum expression within each joint. Using either all markers or differentially expressed markers revealed hand FLS have significantly increased ‘activated’ to ‘resting’ marker ratio^21^. **\*, **, \*\*\*** are p-value < 0.05, 0.01, and 0.005, respectively. The table indicates the number markers with greatest expression in that joint. For example, using all markers, hand FLS has the greatest expression of 66 activated and 42 resting markers. Hand FLS retains the greatest ratio when limiting markers to DEGs. **C**. Heatmap of differentially expressed cytokines and MMPs in TNF stimulated FLS. Rows indicates what joint has the greatest expression of that gene. TNF-stimulated hand FLS have the greatest number of cytokines and MMPs with greatest expression compared to the other joints. **D**. Proliferation and migration biologic validation. Unstimulated and PDGF-stimulated hand FLS proliferate faster than hip or knee. We also evaluated differences in unstimulated and PDGF-stimulated migration and found that hand FLS exhibited greater migration capacity than hip or knee FLS. **\*, **, \*\*\*** are p-value < 0.05, 0.01, and 0.005, respectively.

#### Joint-specific response to TNF reflect differential cytokine and MMP signaling

We then determined the cytokine and MMP expression in TNF stimulated FLS to assess whether those genes reflect the responses associated with the activated fibroblast state. We first evaluated cytokines and MMPs that were differentially expressed between unstimulated vs TNF-stimulated FLS and determined the TNF-stimulated FLS joint with the greatest expression (Figure 4C). Hand and hip FLS had the greatest and least upregulation of cytokines and MMPs, respectively (Figure 4C). The burden of cytokine and MMP gene expression was determined and showed that hand FLS had significantly increased cumulative induction when using either differentially expressed or all cytokines and MMPs (Supplementary Figure 10, Supplementary figure 11) (Permutation test, p-value < 0.05).

#### Increased growth and migration of RA hand FLS

Our computational predictions also suggested that hand FLS might be more aggressive and exhibit differential proliferation and migration compared with hip and knee FLS. Potential differences in proliferation were biologically validated using a cell growth assay, which showed that unstimulated and PDGF-stimulated hand FLS proliferate faster than hip or knee (one-way ANOVA, p-value < 0.05) (Figure 4E). We also biologically tested differences in unstimulated and PDGF-stimulated migration and found that hand FLS migration was greater than hip or knee FLS (Figure 4E).

## Discussion

RA is a symmetric polyarticular arthritis with a characteristic joint distribution. The small joints of the hands, especially the metacarpal-phalangeal joints, are often affected first and can be the most symptomatic. Although there is variability in the clinical course, the disease often progresses to involve larger joints like knees and hips. The mechanisms that explain the distribution of joints in RA are not understood and could relate to the unique biomechanics of each joint^7–9^ or other intrinsic factors. Alternatively, synovial cells might be programmed with joint-specific functions that can participate in embryonic development^22^ as well as contribute to the RA clinical phenotype.

Our epigenetic studies of synovial inflammation in RA have focused on FLS as key stromal elements that can shape the joint structure and function. These cells display a unique aggressive phenotype in RA that contributes to synovial inflammation and joint damage^23,24^. Interestingly, they can potentially spread disease through migration of activated SFs between joints^23^. The role of fibroblasts was also supported by data demonstrating an increase in circulating mesenchymal cells known as PRIME cells prior to RA disease flares^25^. Our understanding of FLS biology was advanced by the discovery of aberrant epigenetic marks involving genes and pathways associated with inflammation and cell migration and invasion in cultured RA FLS ^12,19,20,26^. It is not clear whether these marks are imprinted before cells leave the bone marrow or whether this occurs in situ after cells have migrated to the synovium.

In addition to these overall RA-related epigenomic changes in FLS, marks that are specific to individual joint locations have also been described. Those data largely focused on DNA methylation and identified two types marks: 1) disease-independent epigenetic marks that are found mainly in genes like the homeobox family and likely play a role in embryonic development; and 2) disease-specific marks involving genes and pathways that participate in inflammation and distinguish FLS isolated from different joint locations^12,14^. The lncRNA HOTAIR has also been implicated as one mechanism for joint-specific gene expression^13^. Its effects are limited to a subset of hand-knee differences and are not likely responsible for knee-hip differences. Moreover, our analysis reveals that the regulatory network is likely responsible for the lack of HOTAIR expression in hand FLS and that a much broader regulatory network beyond that lncRNA is defined by the joint of origin. To define these interactions, we generated a new dataset that compares features of resting and TNF-stimulated FLS derived from hands, knees and hips and analyzed it using a novel algorithm that integrates transcriptome and chromatin accessibility. Distinctive joint-specific patterns of TF and regulatees suggest that hand FLS are broadly programmed to be more aggressive than FLS derived knees or hips, including an array of poised enhancers and promoters that can be activated by TNF. These observations, along with biological validation of computational predictions, suggest that FLS imprinting contributes to rheumatoid clinical phenotype.

The Taiji method previously showed that patients could be stratified into biologically distinct groups based on their individual FLS biology and TF function^15^. The method integrates subtle differences between thousands of transcriptional and chromatin accessibility features that are globally summarized by PageRank scores of each TF. Evaluation of differential TFs and regulatees reflect the inherent epigenetic, transcriptomic, and TF differences that synergistically drive joint-specific FLS behavior. As noted above, the method has advantages over analyzing individual datasets like RNAseq or ATACseq. For example, the Taiji networks revealed unique inflammatory (e.g. TLR1:2 Cascades, Interferon alpha/beta signaling), proliferative (e.g. FOXO-mediated transcription of cell cycle genes), and matrix-related (e.g. Laminin interactions) pathways that were not identified with DEGs or DARs alone (Supplementary files 1, 2). This work demonstrates the potential for integrative analyses to identify relevant pathways and therapeutic targets.

Unstimulated cultured RA FLS had transcriptomic and epigenetic patterns that distinguished between joints, as previously reported^12,14^. PCA using either transcriptomic or epigenetic features showed that hand FLS segregate from hip and knee FLS. Likewise, focused unsupervised clustering using only cytokines or limb development factors identified a similar pattern with greatest similarity of transcriptomic patterns shared between hip and knee that was distinct from hand FLS. These observations are consistent with other recent studies where hand and shoulder FLS were distinct from knee^12,14^. Integration of our data by Taiji identified FLS derived from hand had enrichment of proliferative and proinflammatory regulatees of the TFs with the highest PageRank. Epigenetic differences had amplified enrichment of proliferative and inflammatory features in chromatin accessibility that were not yet reflected by the transcriptome, indicating the FLS from the different joints may be epigenetically poised.

Perhaps the most notable results showed differences between joints were amplified with TNF. FLS derived from hands, which already had features that correlate with more aggressive behavior, were distinguished by further enhancement of proinflammatory, proliferative, matrix-destructive genes compared with the other joints. These data are consistent with other recent studies that have shown that hand fibroblasts have greater MMP13 expression after TNF stimulation^12^. Joint-specific sensitivity of FLS and clinical responses in RA to methotrexate and tofacitinib have been observed^18,27^, and it is also possible that responses to TNF inhibitors could be influenced, in part, by joint location. In addition, our study suggests that hand FLS displays the greatest global epigenetic plasticity based on differential regulation of key chromatin remodelers and the greatest global epigenetic changes in response to TNF. It is not certain whether these marks are pre-programmed or whether different mechanical loads modulate distinct inflammatory signaling^28,29^ and epigenetic changes^30^. Thus, the mechanisms of imprinting are largely unknown.

Taiji identified hand FLS are fundamentally distinct from hip and knee in medium and TNF-stimulated conditions. We used the recently reported “activated” and “resting” markers^21,31^ to corroborate our computational predictions in FLS severity and aggressiveness between the joints. We found that unstimulated hand FLS had significant transcriptomic upregulation of “activated” genes and lowest expression of “resting” genes with marked upregulation of cytokines and MMPs in TNF-stimulated hand FLS. This observation agrees with previous work which report greatest expression of cytokines/MMPs in “activated” cells^21^. Unstimulated and TNF-stimulated FLS also had distinct expression and chromatin accessibility profiles of genes and pathways implicated in proliferation and migration. Our biologic validation studies also confirmed that hand FLS had displayed increased proliferation and migration compared with hip and knee FLS.

Our work does have some limitations. First, we used bulk transcriptomic and epigenomic sequencing, which might limit the number of joint-specific fibroblast subpopulations that might drive heterogeneity if studied at a single-cell scale. This present study also does not differentiate joint-specific differences that are disease-independent or disease-dependent due to limitations in sample acquisition. We also cannot differentiate between synovial lining vs sublining, which may be represent distinct cell states^21^. Despite these limitations, we were able to identify significant enrichment of the ‘activated’ state signature recently identified in lining and sublining RA fibroblasts ^21^. Synovial fibroblasts cultured from synovium (conventional FLS) converge to a common phenotype in culture^21,32^, so the relationship between the in vivo and in vivo categories are still uncertain. However, the correlation between the active state and our integrated analysis is striking and suggests that it is relevant to cultured cells.

In conclusion, we performed an integrative genomic analysis that identifies joint-specific differences in FLS biology. The correlation with distribution of joints and the cadence of joints as RA progresses is intriguing and reiterates the importance of joint-context in synovial biology. This, along with other influences like biomechanics, lymphatic distribution and innervation, could contribute to the distribution of joint involvement in RA. Further investigation into features that shape topological potential for inflammation might provide insight into joint-specific pathways that could guide therapy.

## Methods

### Synovial tissue and FLS

Synovial tissue was obtained from RA patients at the time of clinically indicated arthroplasty from 3 joint locations: knee, hip and hand (metacarpal phalangeal joint and wrist). The diagnosis of RA conformed to American College of Rheumatology 2010 criteria^33^. The synovium was processed as previously described, briefly cells were cultured in Dulbecco’s modified Eagle’s medium (DMEM; Life Technologies) supplemented with 10% heat-inactivated fetal calf serum (FCS) (Gemini Bio-Products), and supplements (penicillin, streptomycin, gentamicin, and glutamine) in a humidified atmosphere containing 5% CO2. Cells were allowed to adhere overnight, and then nonadherent cells were removed. Adherent FLS were split at 1:3 when they were 70%–80% confluent and used from passages 4 through 7.^34^ Ten FLS lines each derived from hip, knee and hand synovium were used for subsequent experiments. 70-80% confluent cultures were exposed to media or 50 ng/ml of TNF for 6 hours the cells were harvested and processed for ATACseq or RNAseq.

### RNA-seq and ATAC-seq sample processing and assays

Genomic DNA and total RNA from parallel FLS cultures were used for RNAseq and ATACseq. Total RNA was extracted and the quality of all samples was evaluated using an Agilent Bioanalyzer. The samples had an average RNA Integrity Number (RIN) of 9.4 with a minimum of 8. Sequencing libraries were prepared using TruSeq Stranded Total RNA RiboZero protocol from Illumina. Libraries were pooled and sequenced with an Illumina HiSeq 2000 (Institute for Genomic Medicine, UCSD). Genomic DNA was processed for ATACseq (Center for Epigenomics, UCSD) and subsequently sequenced (Institute for Genomic Medicine, UCSD)

### RNA-seq and ATAC-seq data processing

To facilitate reproducible results, RNA-seq and ATAC-seq fastq files were processed using the nf-core pipeline<u>^35^</u>, which verifies raw sequencing quality, trims adapters, removes genome contaminants, removes ribosomal RNA, deduplicates, and conducts additional extensive quality control for RNA-seq. For ATAC-seq, nf-core pipline verifies raw sequencing quality, trims adapters, deduplicates, removes mitochondrial DNA, blacklisted regions, low-quality unmapped, multi-mapped, mismatched, or multi-chromosomal-matched alignments, and conducts additional extensive quality control. No data were excluded from RNA-seq or ATAC-seq analysis for quality reasons. Environment for nf-core/RNAseq and nf-core/ATACseq are as follows: Singularity v3.8.6-1.e18, nextflow v22.04.0, with either nf-core/rnaseq v3.8.1 or nf-core/atacseq v1.2.1 with all default nf-core parameters. Briefly regarding the RNA-seq processing workflow, reads were trimmed with Trim Galore! v.0.6.7, aligned to hg38 with STAR v2.7.10a, and bam-level quantification performed with salmon v0.13.1. For ATAC-seq, reads were trimmed (as above), aligned to hg38 with BWA v0.7.17, blacklisted regions^36^ were filtered, and narrow peaks were called with macs2 v2.2.7.1. Only autosomes were used for downstream analysis. RNA-seq was sequenced with only one batch. ATAC-seq was sequenced with two batches. No batch effects were observed (Supplementary figure 12). All differentially expressed genes (DEGs) were found with the nonparametric Wilcoxon test (p-value < 0.05, log2FC > 0.58). All differentially accessible region (DAR) analysis (FDR < 0.05, log2FC > 0.58) was conducted with the DiffBind^37^ R package.

### Construction of genetic networks using Taiji

We used the Taiji^15,16^algorithm (v.1.3.0) to incorporate RNA-seq, ATAC-seq, and enhancer-promoter interactions (the top 10% most confident predictions from Epitensor^38^) (Supplementary figure 13). For each sample, Taiji produced a genetic network and inferred TF importance via personalized PageRank calculations for 745 TFs with known motifs. Briefly, the nodes in the network are genes and are weighted by the normalized gene-expression level. To create edges between regulators and regulatees, open chromatin promoter and enhancer regions are queried for TF motifs documented in the Cis-BP^39^ database. These edges are weighted by the motif score reflecting the binding affinity, TF expression levels, and target open chromatin peak intensity. The personalized PageRank algorithm is then applied to the resulting directional and weighted network to determine relative importance of each TF for each sample. The output of the Taiji workflow for each sample are the personalized PageRank scores for 745 TFs, the network topology consisting of directional regulator—regulatee relations with each pair’s associated edge weight and node weights. Individual genetic networks were constructed for 30 RA FLS cell-lines derived from various anatomical locations (hand, knee, hip) and conditions (unstimulated, TNF) for 60 total samples. Most confident regulator-regulatees edges (shared by >70% samples within joint and condition) were used for downstream analysis (Supplementary file 6).

### Proliferation and migration assays

For proliferation, FLS were plated in 96 well plates, 3000 cells/wells in quadruplicates as described. Cells were serum starved with 1%FBS for 24h followed by incubation with 1%FBS with and without PDGF (10ng/ml). 20 ul of MTT (Invitrogen) were added in each well for and the plates read at 550nm using 690nm as reference. For migration, RA FLS were plated into 12 well plate (1.5×10^5^ cell/well) and starved with 0.1% FCS DMEM and a scratch introduced with a pipette tip. After culturing for 24 h in the presence or absence of PDGF (10 ng/ml) in 0.1% FCS DMEM, cells are fixed with 4% PFA and stained with crystal violet. They were visualized using an ECLIPSE E800 (Nikon) microscope at 40x magnification and counted with ImageJ distance. Cell proliferation was performed as previously described^40^.

### Permutation tests

Significant enrichment of activation markers between joints *i* and *j* are identified using a permutation test by randomly shuffling the joint labels with the greatest expression (defined by the median for each joint) of that activated/resting marker and then recalculating the ratio of max #activated to #resting markers. The test statistic is defined as:

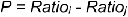

We also identified enrichment of cumulative cytokine expression in TNF-stimulated cell-lines by taking the cumulative expression of cytokines and MMPs averaged (by median) per joint. For each pairwise comparison, significant upregulation of cytokines between joints *i* and *j* are identified using a permutation test by randomly shuffling the joint labels and taking the sum expression:

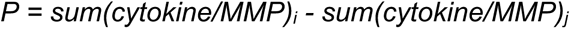

For both permutation tests, the p-value of is computed by:

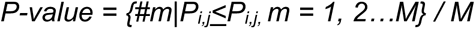

with M = 5000 permutations.

### Study approval

These studies were approved by the UC San Diego Institutional Review Board (IRB #014-175). All research participants signed informed consent.

### Data availability

The data that support the findings of this study are available in dbGaP (accession #xxxxxx).

## Supporting information

Supp 4 diffbind outputs

Supp 6 confident edges

supp 5 differential PageRanks

supp 3 DEG analysis

Supp 2 TNF pathways

supp 1 control pathways

## Acknowledgements

Supported by grants AR071321 and AR065466 from the National Institutes of Arthritis and Musculoskeletal and Skin Diseases

---

**S1.**
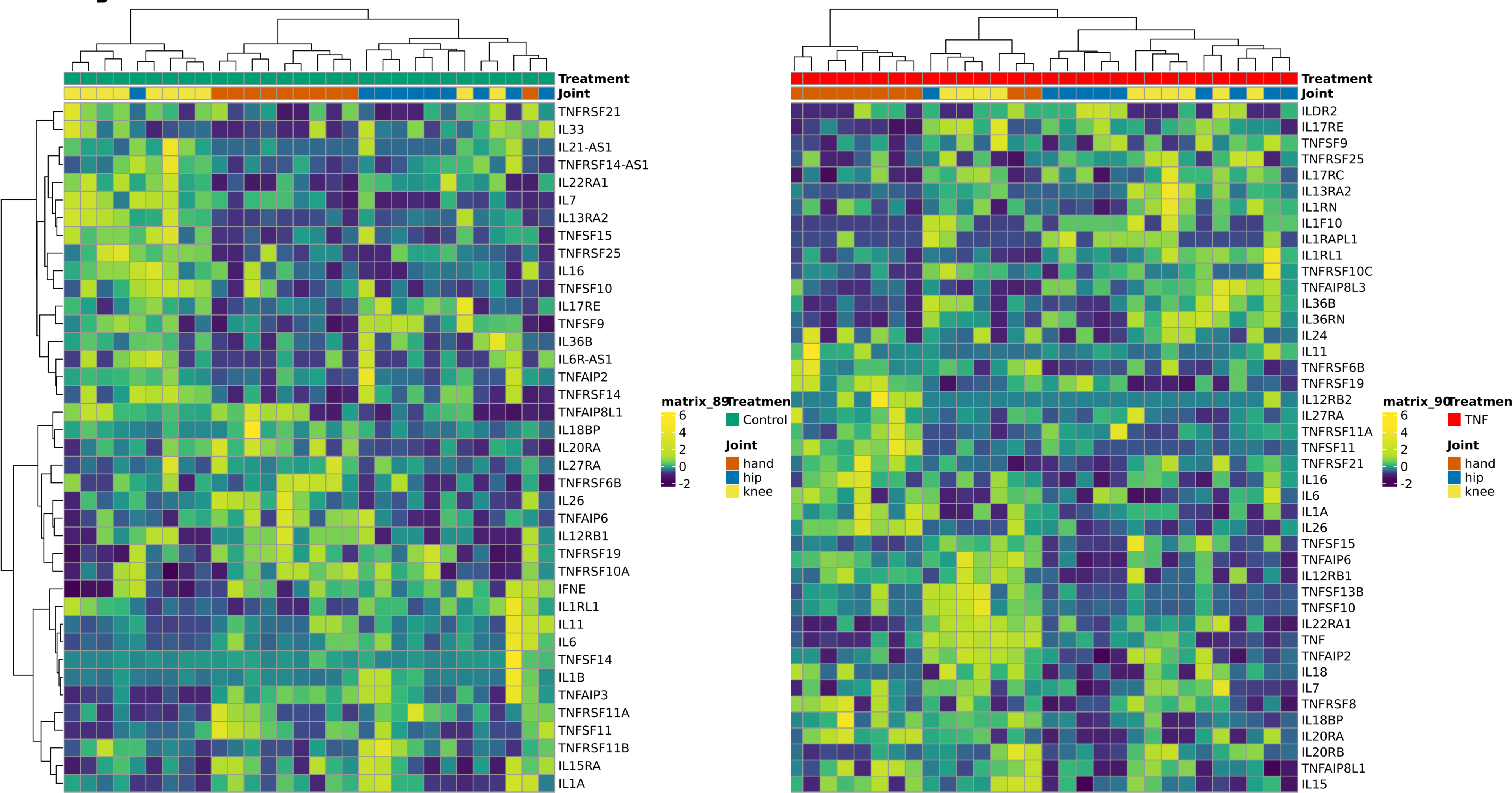
Unsupervised clustering using differentially expressed cytokines. Heatmap using unsupervised clustering of differentially expressed cytokines within medium (left) or TNF-stimulated (right) conditions. Ward’s Hierarchical Agglomerative Clustering Method using the correlation distance was used to cluster samples by cytokine expression. Unstimulated (left) and TNF-stimulated (right) FLS exhibit the greatest mixing with hip and knee.

**S2.**
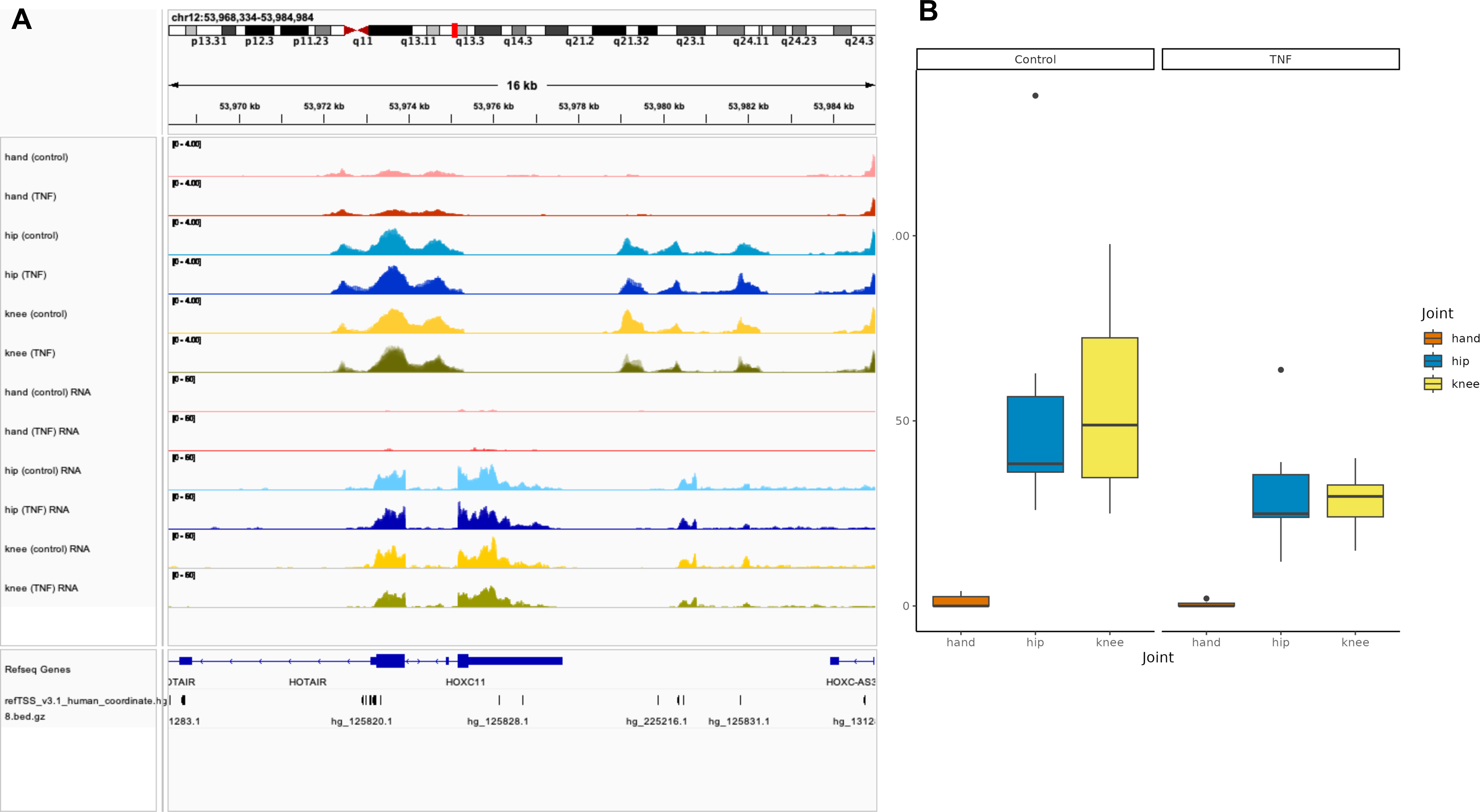
HOTAIR chromatin accessibility and gene expression plots. Genome browser of chromatin accessibility (top 6, ordered by joint and treatment) and gene expression (bottom 6, ordered by joint and treatment). View **S3** for significance values. **B.** Box plot of gene expression profiles of HOTAIR within treatment and between joints. View **S4** for significance values.

**S3.**
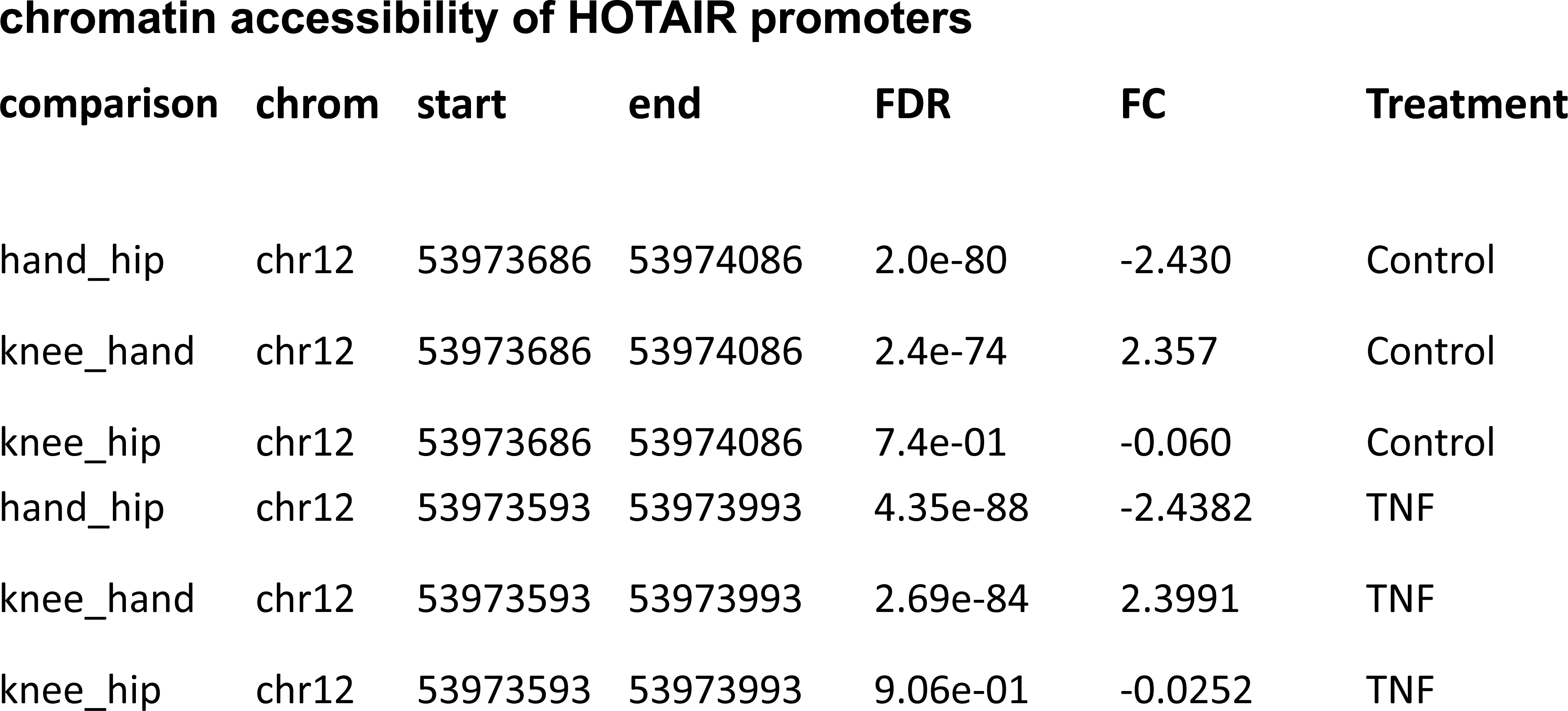
HOTAIR chromatin accessibility tables. HOTAIR promoter chromatin accessibility significance values within treatment and between joints. FC is first joint / second joint. For example, TNF knee_hand has FC of 2.4. This means knee has a higher peak than hand by 2.4.

**S4.**
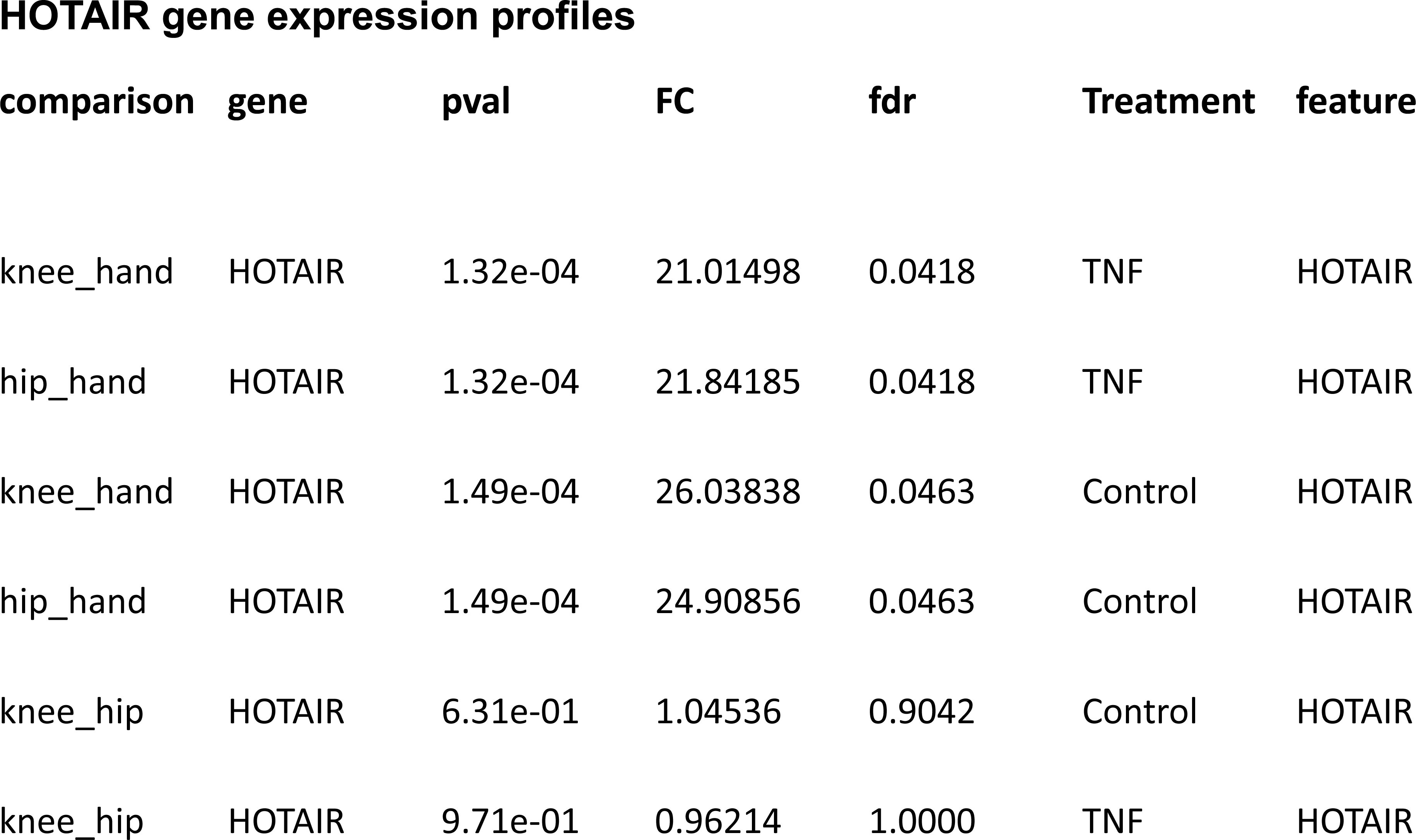
HOTAIR gene expression tables. HOTAIR gene expression significance values. FC is first joint / second joint. For example, TNF knee_hand has FC of 21.01. This means knee has higher HOTAIR expression than hand by 21.01.

**S5.**
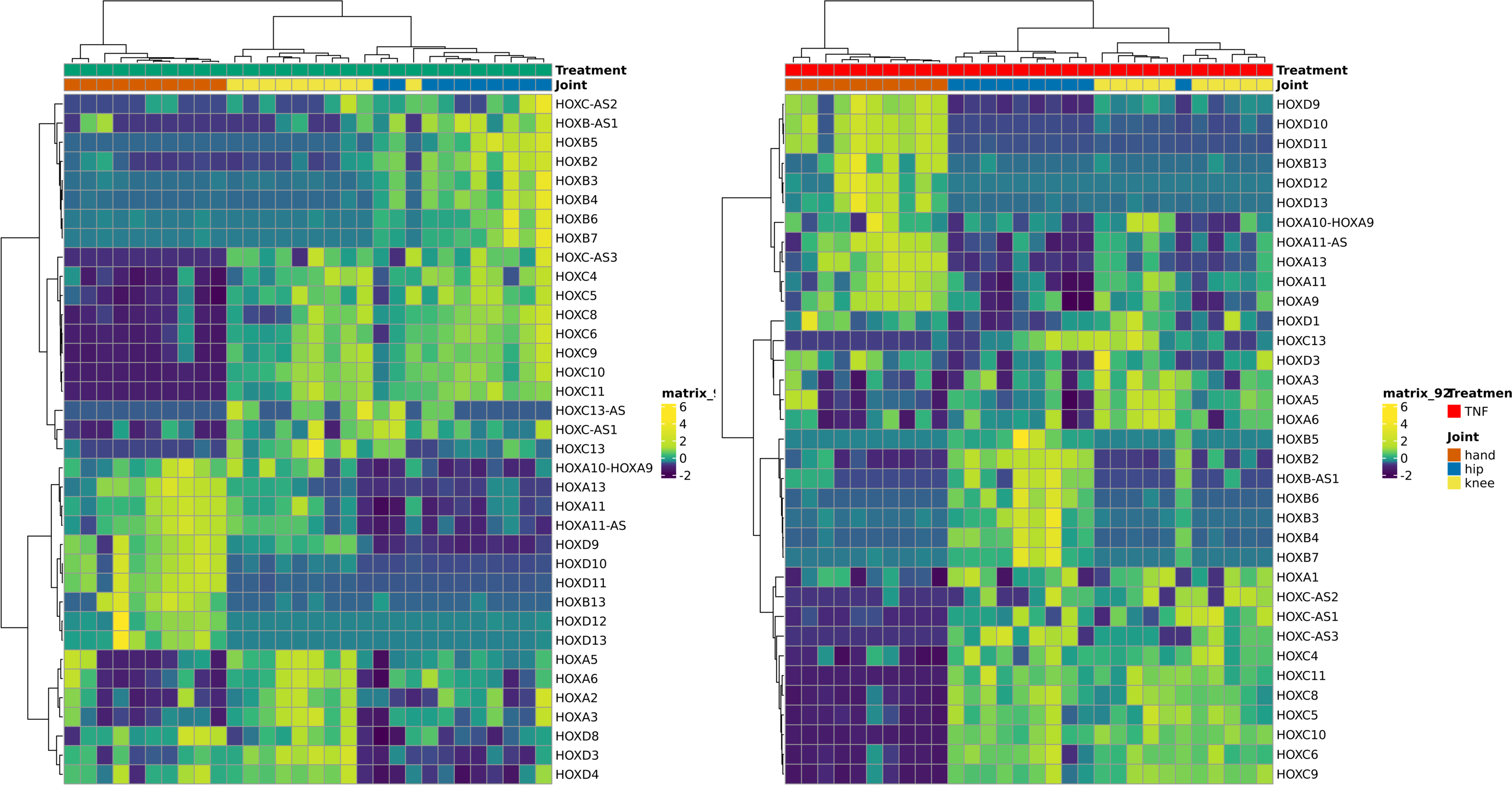
Unsupervised clustering using differentially expressed limb patterning genes. Hierarchical clustering within unstimulated or TNF-stimulated FLS (ward.D2, correlation method). Heatmap using unsupervised clustering of differentially expressed HOX genes within unstimulated (left) or TNF-stimulated (right) conditions. Consistent expression patterns between hip with knee were observed whereas hand FLS had the most distinct expression compared with the other joints.

**S6.**
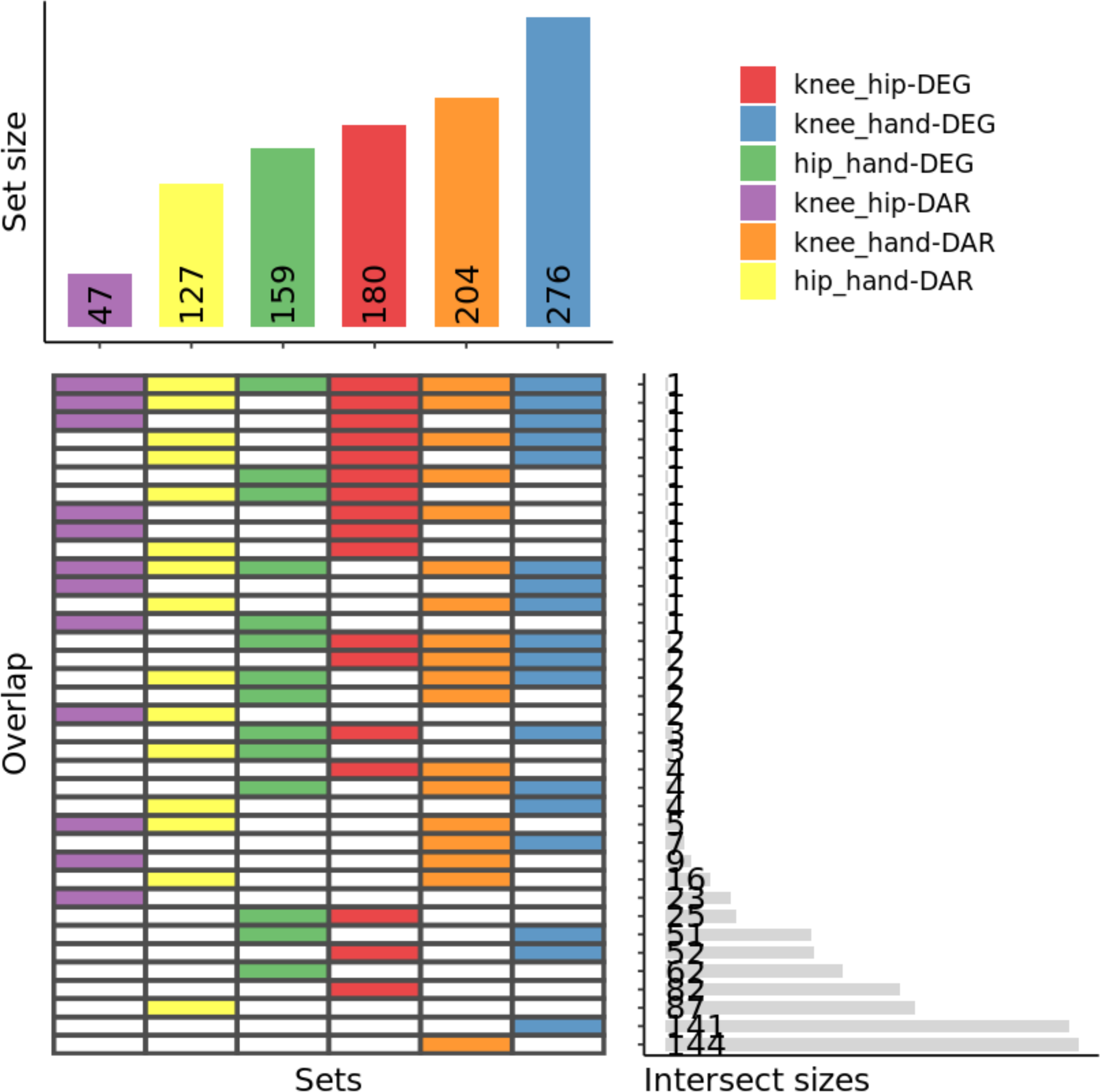
Unstimulated FLS have little overlap between DEGs and DARs. Upset plot that visualizes the overlap of DARs and DEGs for each pairwise comparison within unstimulated FLS and between hand, hip, and knee. This is read identical to Figure 2B and Figure 3B. The top bars indicate number of all features for that group. The horizontal bars indicate the number of features within that overlap subsection. For example, hand vs hip DARs (yellow) and DEG (green) only have 3 overlapping features indicating FLS are in an epigenetically poised state.

**S7.**
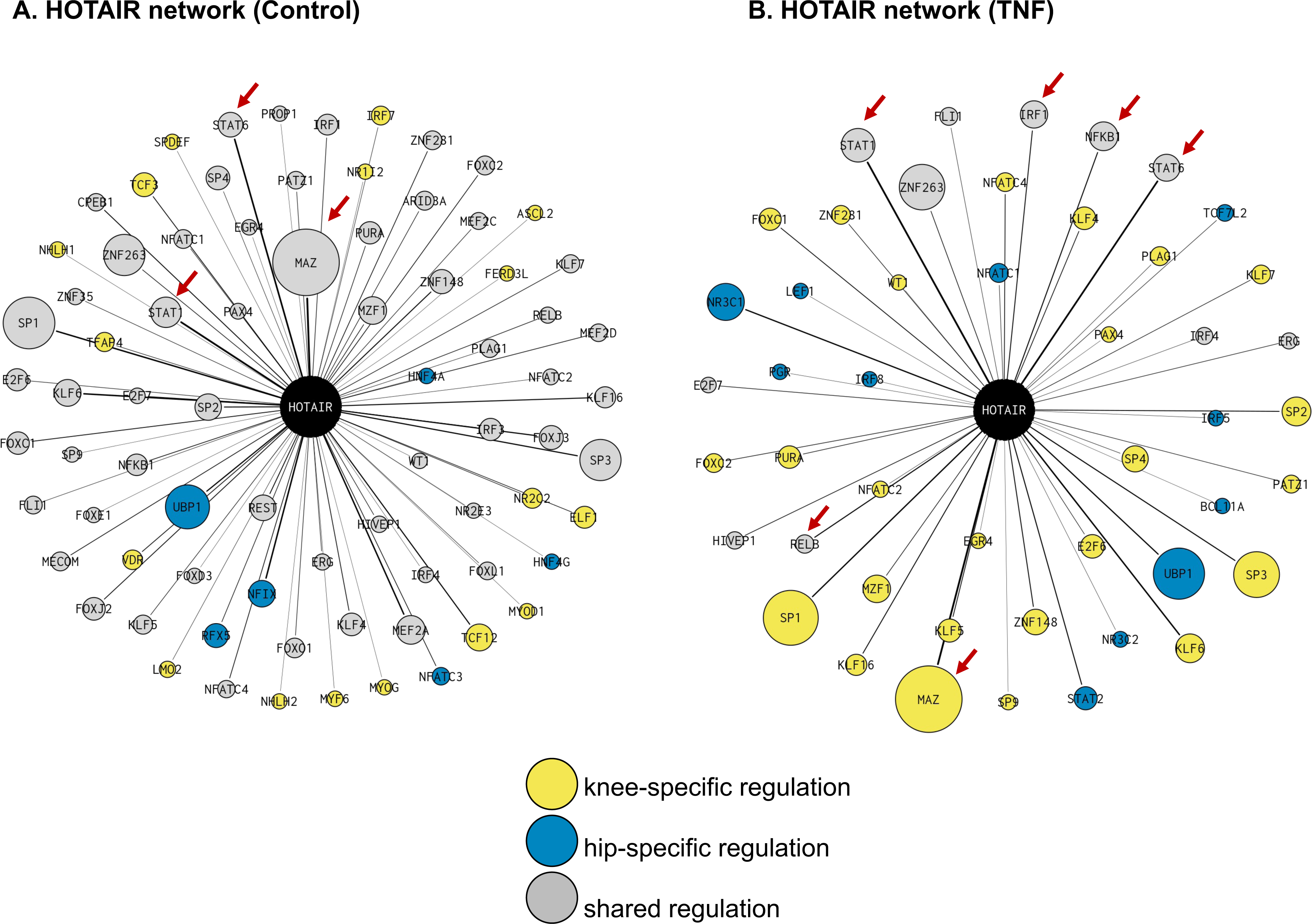
HOTAIR transcriptional network identifed from Taiji. For each treatment (Control, TNF), we identified joint-specific and shared regulation of HOTAIR. Nodes (circles) are scaled by PageRank (higher PageRank → bigger node). Nodes are colored blue (hip-specific), yellow (knee-specific), or grey if the regulatory relationship is shared by both hip and knee. If the node is joint-specific (i.e. blue or yellow color), the PageRank size is scaled by average PageRank of that joint. If the node is shared (i.e. grey), the PageRank size is scaled by the average PageRank across knee and hip. Edges are weighed by the average edge weight (same method as node weight). The darker the edge, the higher the edge weight and thus the greater the regulatory potential between that TF to HOTAIR. Key TFs are indicated with red arrows.

**S8.**
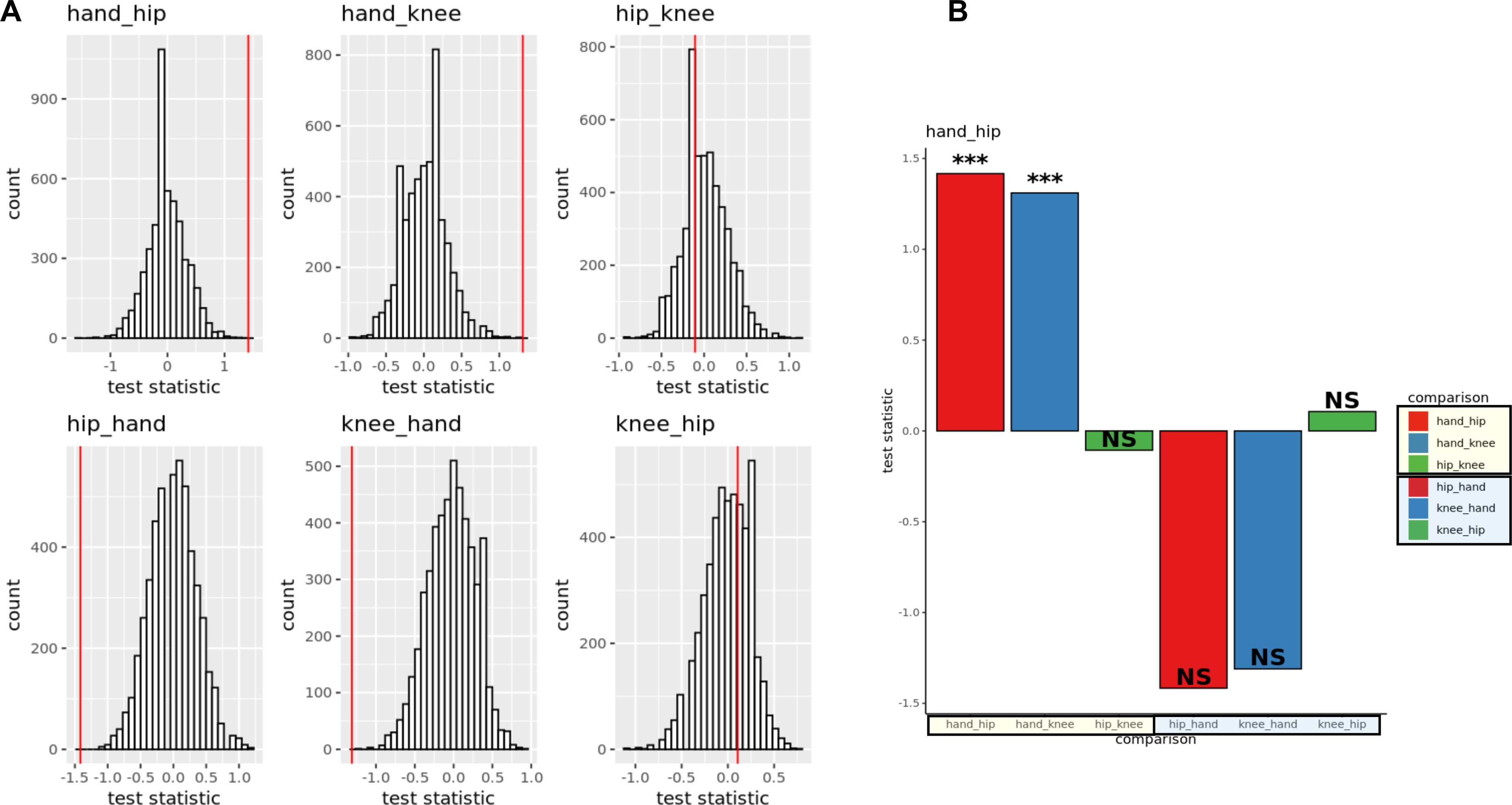
Hand FLS has significant enrichment of genes associated with the ‘activated’ state (DEGs). Significant enrichment of activation markers between joints *i* and *j* are identified using a permutation test by randomly shuffling the joint labels with the greatest expression (defined by the median for each joint) of that activated/resting marker and then recalculating the ratio of max #activated to #resting markers. The test statistic is defined as: 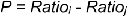 The p-value of is computed by: = *P-value = {#m|P_i,j_<P_i,j,_ m = 1, 2…M} / M* with M = 5000 permutations. Figure **S8A** is the histogram of test statistics of shuffled activated/resting marker labels. The vertical line is the original test statistic. For example, in hand_hip plot, we hypothesized that hand would have a greater ratio of activated to resting marker expression. The original test statistic > 1 indicating hand FLS has greater enrichment of activated markers compared to hip. P-value < 0.05 indicates this is unlikely due to chance, as shown in S8B. Though knee has the greatest expression of the most activated markers, it is attenuated by increased expression of resting markers. *, **, *** represents p-value < 0.05, 0.01, and 0.005, respectively.

**S9.**
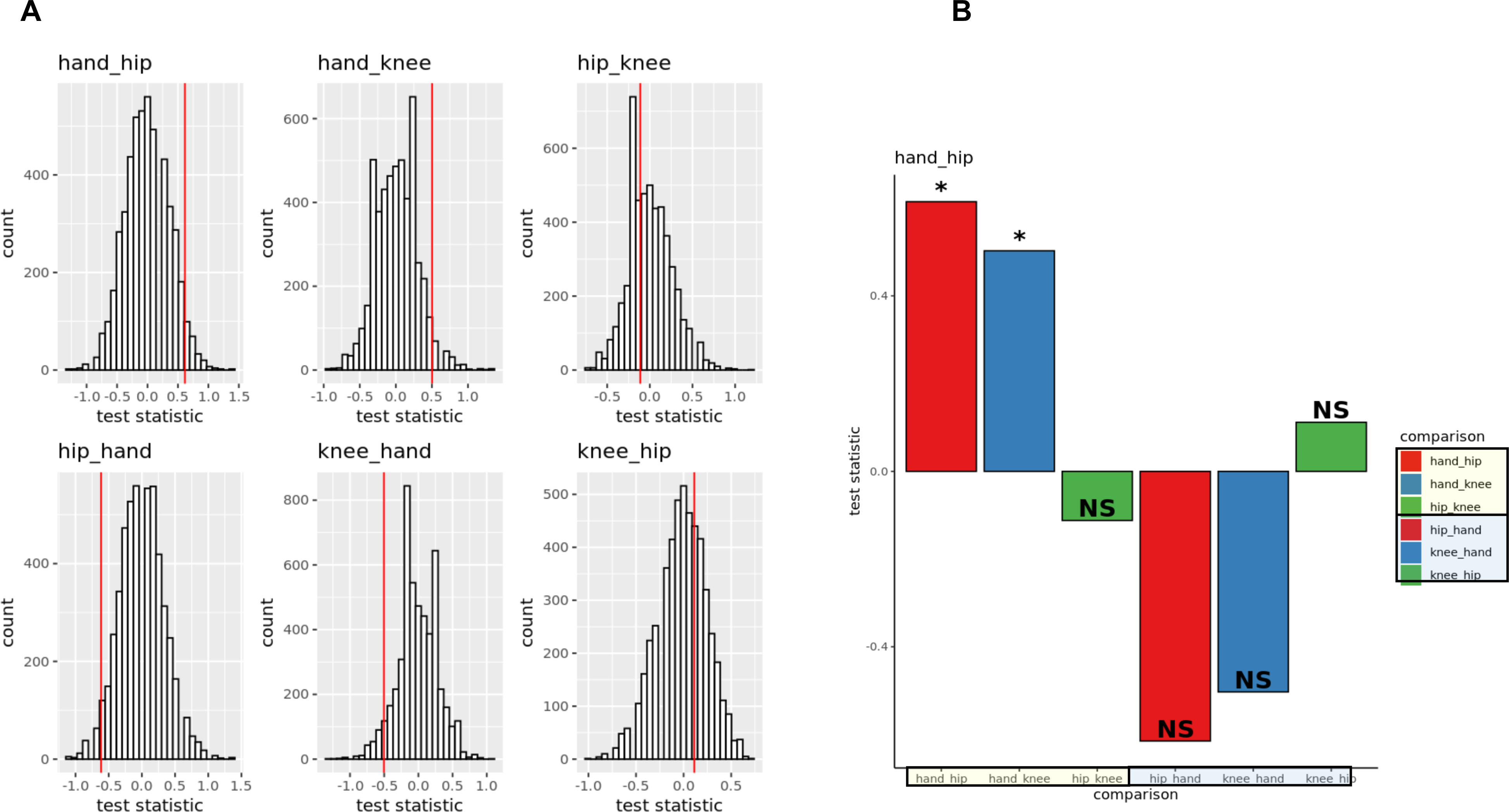
Hand FLS has significant enrichment of genes associated with the ‘activated’ state (all markers). Activated and resting marker expression were evaluated using all markers within unstimulated FLS. Similar observation using differentially expressed markers are observed when using all markers with the greatest activated marker expression observed in hand FLS. **\*, **, \*\*\*** represents p-value < 0.05, 0.01, and 0.005, respectively. See **S8** for more detailed description.

**S10.**
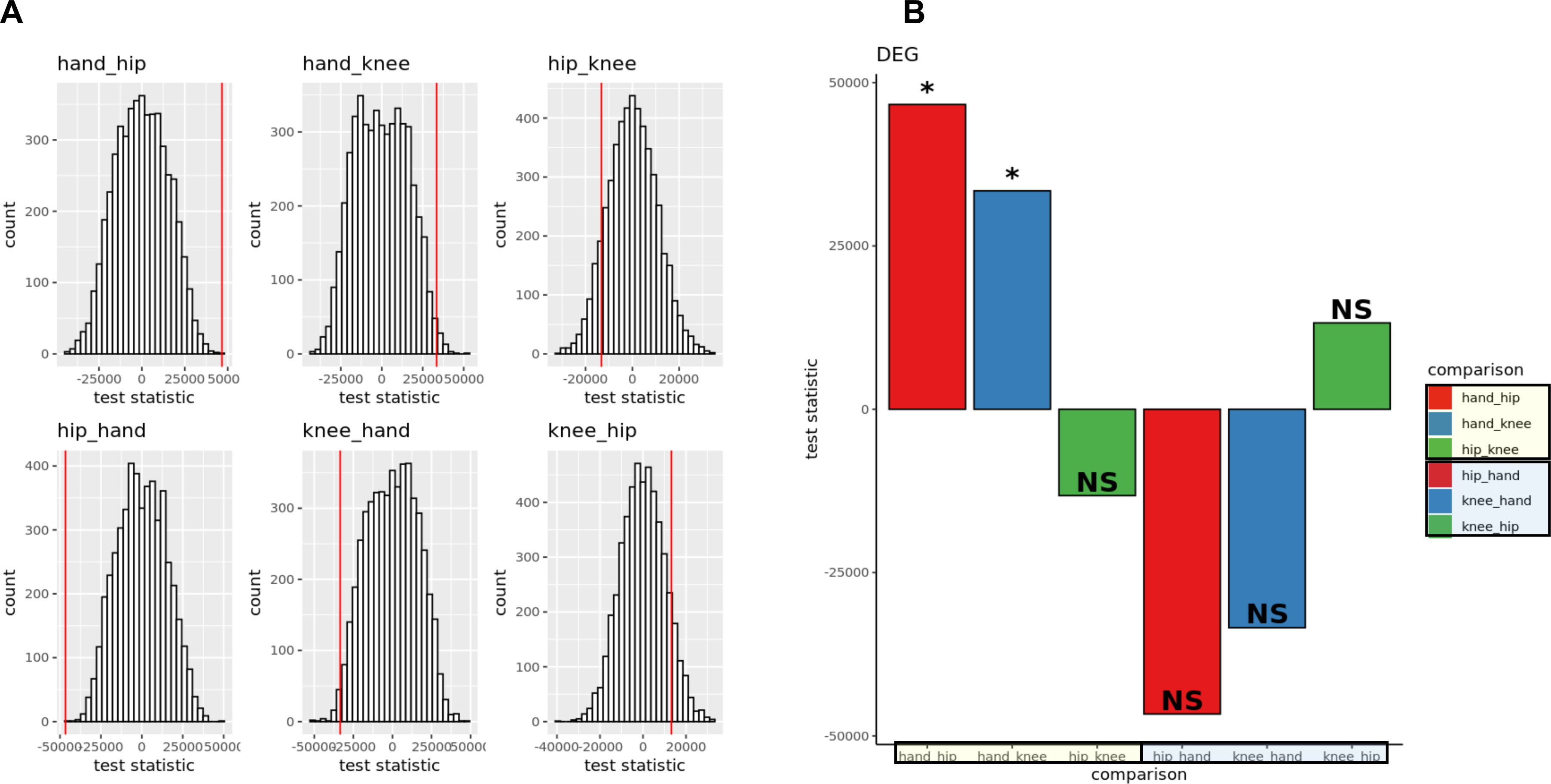
TNF-stimulated hand FLS has increased burden of cytokine and MMP expression (DEGs). We identified the enrichment of cumulative cytokine and MMP expression in TNF-stimulated cell-lines by taking the cumulative expression of cytokines and MMPs averaged (by median) per joint. For each pairwise comparison, significant upregulation of cytokines between joints *i* and *j* are identified using a permutation test by randomly shuffling the joint labels and taking the sum expression: 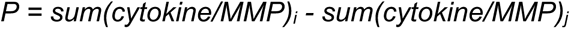 The p-value of is computed by: = *P-value = {#m|P_i,j_≤P_i,j,_ m = 1, 2…M} / M* with M = 5000 permutations. Figure S7.1 is the histogram of test statistics of shuffled joint labels within TNF-stimulated conditions given differentially expressed cytokines. The vertical line is the original test statistic. For example, in hand_hip plot, we hypothesized that hand would have a greatest cumulative expression of cytokines compared to hip. The original test statistic > 1 indicating hand FLS has greatest cumulative expression of cytokines. P-value < 0.05 indicates this is likely not due to chance. *, **, *** represents p-value < 0.05, 0.01, and 0.005, respectively.

**S11.**
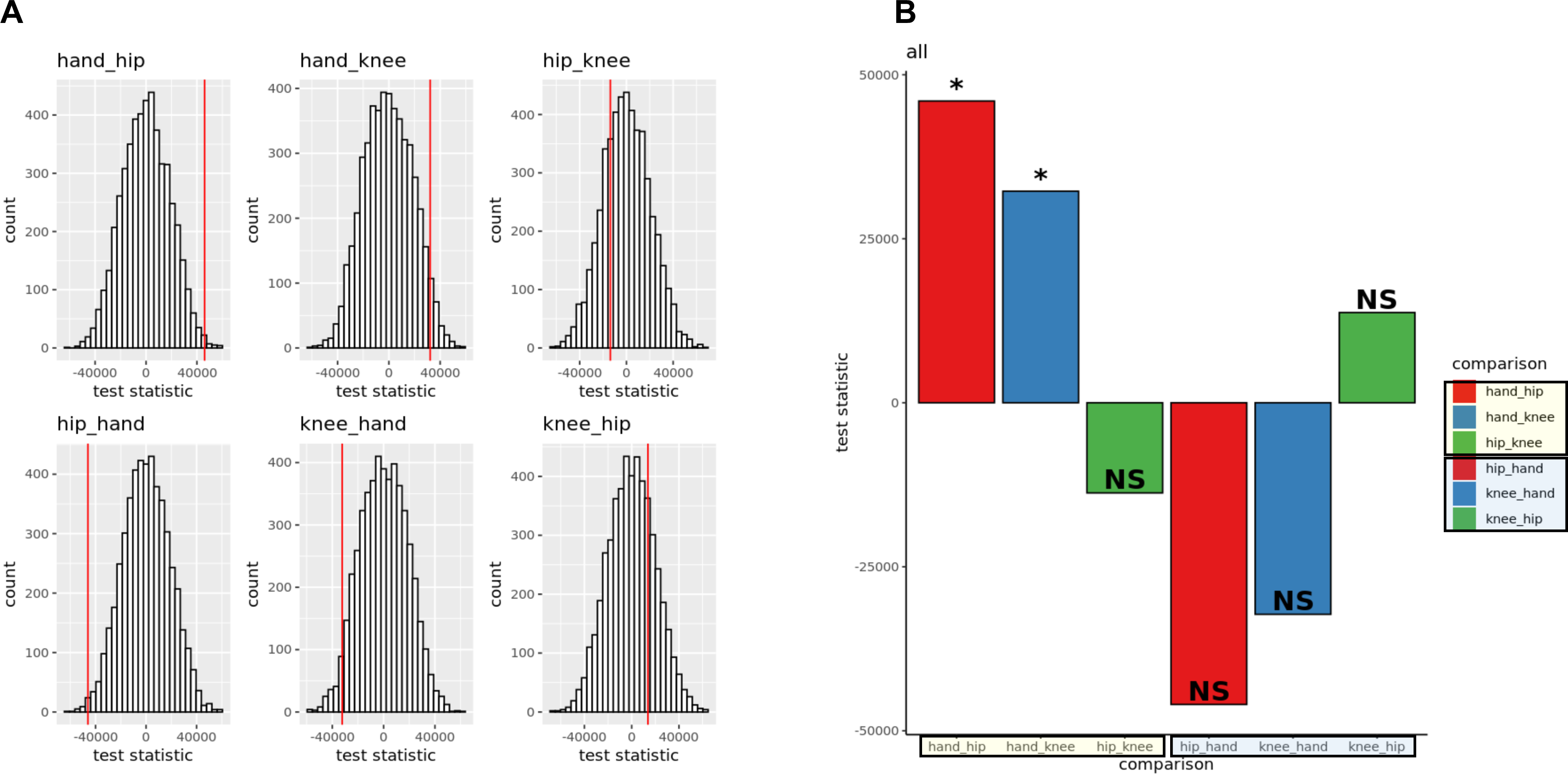
TNF-stimulated hand FLS has increased burden of cytokine and MMP expression (all markers). Cumulative cytokine and MMP expression were evaluated using all cytokines within unstimulated FLS. Similar observation using differentially expressed cytokines are observed when using all markers with the greatest cumulative expression observed in hand. *, **, *** represents p-value < 0.05, 0.01, and 0.005, respectively. View S10 for a more detailed description.

**S12.**
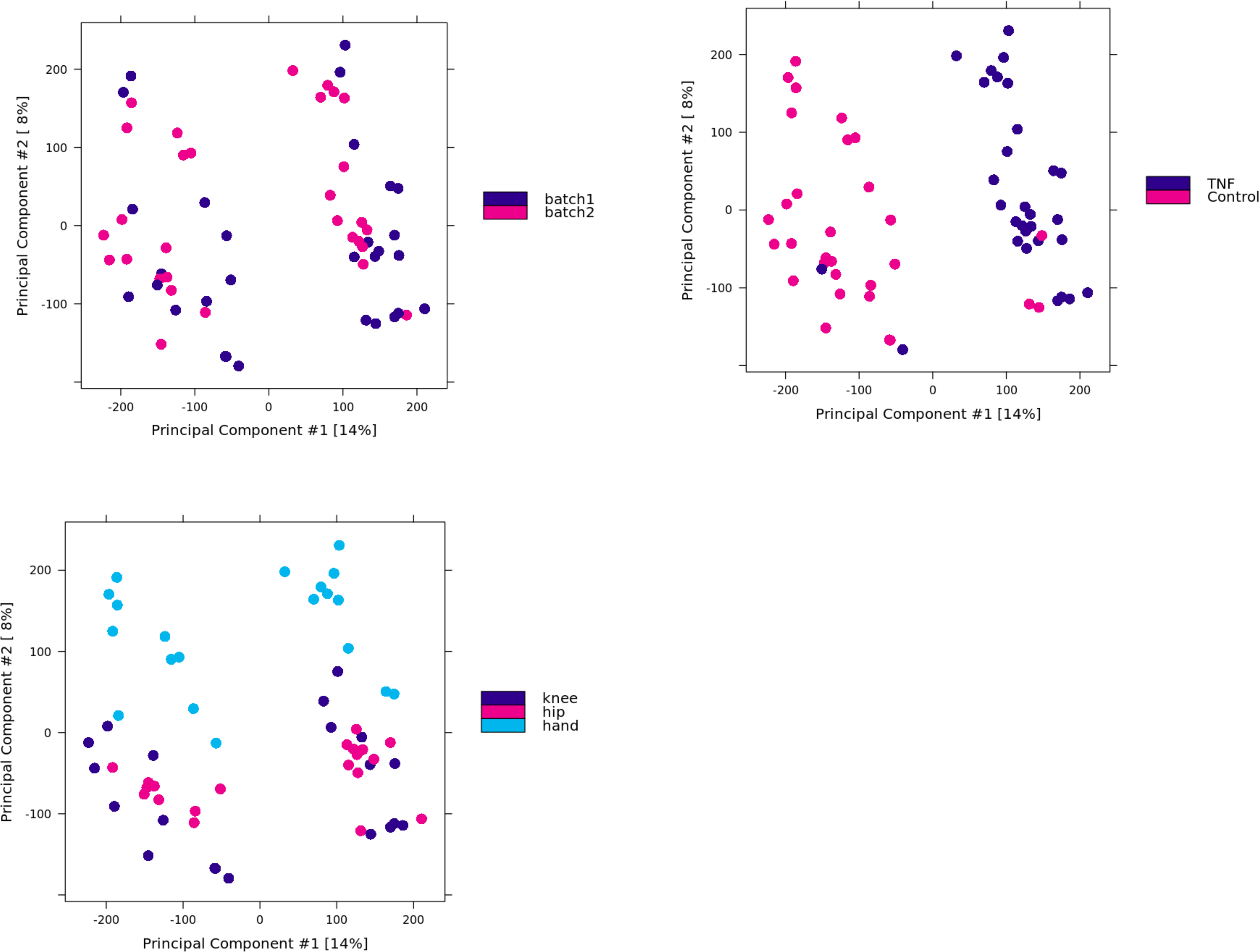
Batch effects are not observed in ATACseq experiments. ATAC-seq files were processed in two batches. Batch effects were evaluated using PCA. Samples segregate by condition (control, TNF) and joint (hand, hip, knee). Batch effects were not observed. NOTE: RNA-seq files were processed in the same batch.

**S13.**
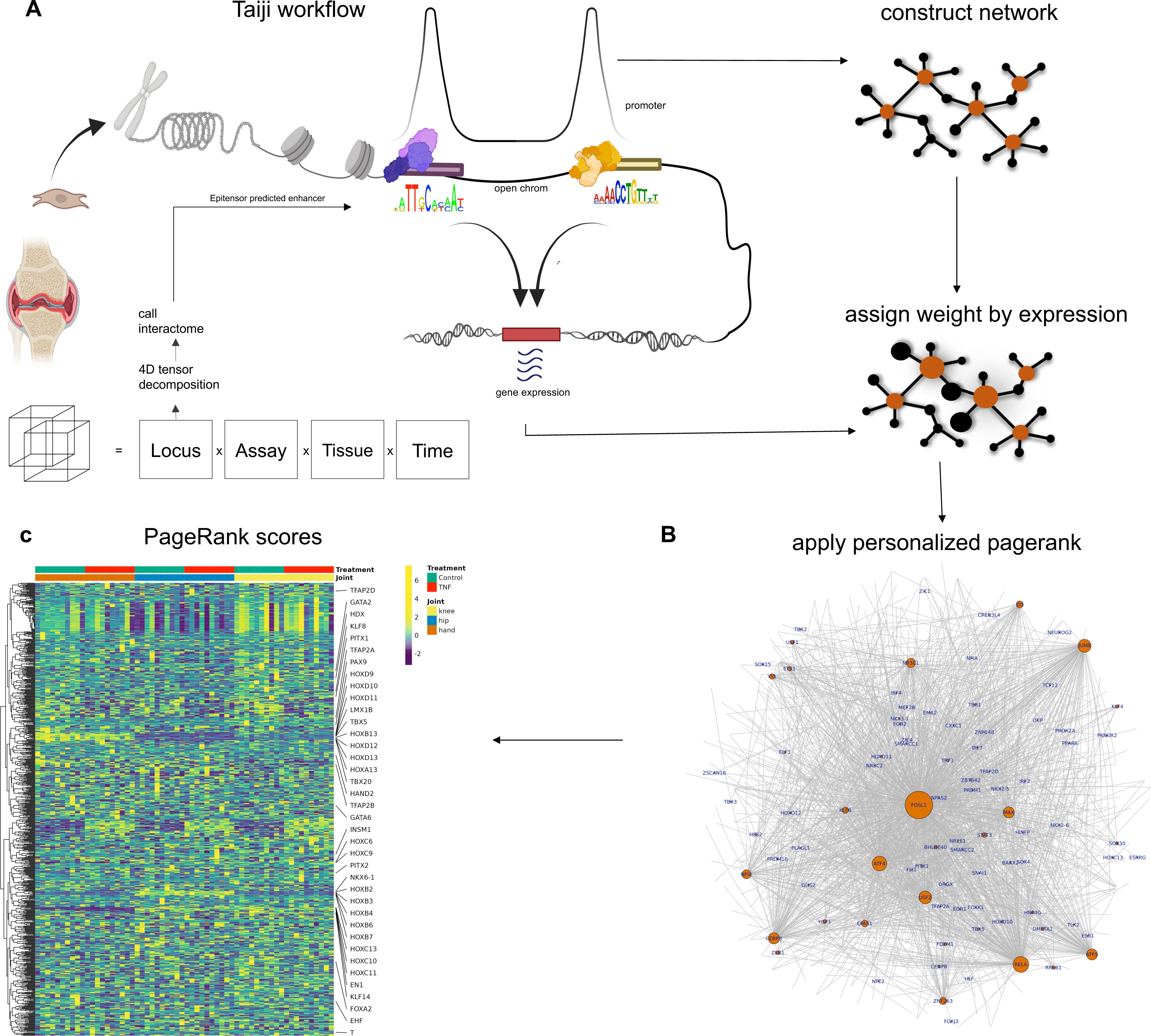
Taiji workflow. Schematic of the taiji workflow. Taiji integrates RNA-seq, ATAC-seq, and enhancer-promoter interactions (the top 10% most confident predictions from Epitensor^8^). For each sample, Taiji produced a genetic network and inferred TF importance via personalized PageRank calculations for 745 TFs with known motifs. Briefly, the nodes in the network are genes and are weighted by the normalized gene-expression level. To create edges between regulators and regulatees, open chromatin promoter and enhancer regions are queried for TF motifs documented in the Cis-BP^9^ database. These edges are weighted by the motif score reflecting the binding affinity, TF expression levels, and target open chromatin peak intensity. The personalized PageRank algorithm is then applied to the resulting directional and weighted network to determine relative importance of each TF for each sample. The output of the Taiji workflow for each sample are the personalized PageRank scores for 745 TFs, the network topology consisting of directional regulator—regulatee relations with each pair’s associated edge weight and node weights.

